# Generalists in a Specialist World: Vibrio-Phages with Broad Host Range

**DOI:** 10.1101/2024.12.24.630238

**Authors:** Charles Bernard, Yannick Labreuche, Carine Diarra, Pauline Daszkowski, Karine Cahier, David Goudenège, Martin G. Lamarche, Gregory Whitfield, Manon Lang, Jeffrey Valencia, Justine Groseille, Damien Piel, Yves V. Brun, Eduardo P.C. Rocha, Frédérique Le Roux

**Affiliations:** Institut Pasteur, Université de Paris, CNRS UMR3525, Microbial Evolutionary Genomics, Paris, France; Ifremer, Unité Physiologie Fonctionnelle des Organismes Marins, ZI de la Pointe du Diable, CS 10070, F-29280 Plouzané, France; Sorbonne Universités, UPMC Paris 06, CNRS, UMR 8227, Integrative Biology of Marine Models, Station Biologique de Roscoff, CS 90074, F-29688 Roscoff cedex, France; Département de microbiologie, infectiologie et immunologie, Université de Montréal, Montréal, Canada

## Abstract

The host range of a bacteriophage—the diversity of hosts it can infect—is central to understanding phage ecology and applications. While most well-characterized phages have narrow host ranges, broad-host-range phages represent an intriguing component of marine ecosystems. The genetic and evolutionary mechanisms driving their generalism remain poorly understood. In this study, we analyzed Schizotequatroviruses and their *Vibrio crassostreae* hosts, collected from an oyster farm. Schizotequatroviruses exhibit broad host ranges, large genomes (~252 kbp) encoding 26 tRNAs, and conserved genomic organization interspersed with recombination hotspots. These recombination events, particularly in regions encoding receptor-binding proteins and antidefense systems, highlight their adaptability to host resistance. Notably, some lineages demonstrated receptor-switching between OmpK and LamB, showcasing their evolutionary flexibility. Despite their broad host range, Schizotequatroviruses were rare in the environment. Their scarcity could not be attributed to burst size, which was comparable to other phages *in vitro*, but may result from ecological constraints or fitness trade-offs, such as their preference for targeting generalist vibrios in seawater rather than the patho-phylotypes selected in oyster farms. Our findings clarify the genetic and ecological trade-offs shaping Schizotequatrovirus generalism and provide a foundation for future phage applications in aquaculture and beyond.

## INTRODUCTION

Bacteriophages (or phages)—viruses that infect bacteria—are attracting renewed attention due to their ecological significance, impact on human health, and potential applications (1–3). Effective therapeutic use of phages requires a host range that encompasses the genetic diversity of bacterial pathogens (4). Host range is primarily determined by the phage’s ability to bind specific bacterial receptors, with variations in receptor-binding proteins broadening or narrowing this range (5, 6). Bacterial defense mechanisms (e.g. (7)) also influence phage infection, and phages evolve rapidly to counter these defenses, with environmental context shaping the co-evolutionary dynamics between phages and bacteria (8). Phages exhibit a wide range of host specificities, from those with highly restricted host ranges to others with broad infectivity, though most well-characterized model phages tend to have narrow host ranges (5). Studies of broad-host-range marine phages and metagenomic analyses of diverse environments suggest that such phages may be more common than previously thought in natural ecosystems (9–11). Yet, little is known about the genetic determinants, evolutionary dynamics, trade-offs, and factors shaping host tropism that underlie the generalism of these phages in natural ecosystems.

Our previous research showed that juvenile oysters impacted by Pacific oyster mortality syndrome (12) are infected by diverse virulent genotypes within the *Vibrio crassostreae* species (13, 14). These virulent strains are largely organized into distinct phylogenetic clades (15), or ‘phylo-pathotypes,’ and carry the virulence plasmid pGV, which encodes a T6SS responsible for cytotoxicity (16). We also found this *V. crassostreae* population to be highly dynamic (13, 15), sparking our interest in the role of phages in shaping its abundance and diversity. To investigate, we collected marine phages and *V. crassostreae* hosts over a time series from an oyster farm and assessed their host range through cross-infection studies (15). This analysis revealed a highly modular infection network, where phage adsorption is governed by specific matches between phage genus and vibrio clades. Following adsorption, intracellular defense systems further narrow the range of strains that are successfully infected and lysed.

One phage in our study, 6E351A, piqued our curiosity as the only isolate among 242 (0.4%) capable of infecting not only *V. crassostreae* strains outside any specific clade, but also strains from other *Vibrio* species (15). Genome sequencing identified this phage as part of the Schizotequatrovirus family (previously known as giant vibriophages or T4 giant phages). These viruses, exhibiting a Myoviridae morphotype, are characterized by large capsids and double-stranded DNA genomes of approximately 250 kbp, with a quarter of their coding regions showing homology to T4 phage genes (17). The best-known member of this group, KVP40, is notable for its broad host range (18). This finding suggests that Schizotequatroviruses may generally include broad-host-range phages. However, research on these phages is limited by the small number of isolated phages and hosts and the considerable genetic diversity within this family.

In this study, we established a collection of Schizotequatroviruses and their *V. crassostreae* hosts to investigate the fine-scale diversity and evolutionary dynamics within this family. Uncovering the genetic mechanisms and evolutionary processes that enable these phages to maintain a broad host range, along with understanding the potential trade-offs that may explain their low prevalence, will provide valuable insights into their role in shaping bacterial eco-evolutionary dynamics and inform their potential applications.

## MATERIAL AND METHODS

### Isolation and identification of Schizotequatroviruses

Sampling was conducted from an oyster farm in the Bay of Brest (Pointe du Château, 48° 20′ 06.19′′ N, 4° 19′ 06.37′′ W) between June 28 and September 15, 2021, on Mondays, Wednesdays, and Fridays. Specific Pathogen-Free juvenile oysters were deployed in consecutive batches, to monitor mortality linked to seawater temperature increases above 16°C, a threshold for oyster mortalities. Samples were collected on each sampling day from living oysters within a batch showing <50% mortality. Hemolymph was collected from 90 living oysters, centrifuged (10 min,17,000 g) and the supernatant filtered through a 0.2 μm filter and stored at 4°C until the phage isolation stage. Concurrently, 10 liters of seawater were collected and size fractionated as previously described (13) and viral particles were concentrated using 0.2 µm filtration and iron chloride flocculation, respectively, following standardized protocols (19). Virus-flocculates were suspended in 10mL 0.1M EDTA, 0.2M MgCl_2_, 0.2M oxalate buffer at pH6 and stored at 4°C until the phage isolation stage.

A total of 153 *V. crassostreae* isolates (15) were used as bait to isolate phages across 35 sampling dates. For each date, a mixture of 10 µL seawater viral concentrate (equivalent to 10 mL of seawater, concentrated 1,000-fold) and 10 µL of oyster plasma derived from a pooled sample of 90 oysters was tested. Phage infections were identified by the formation of plaques using soft agar overlays on bacterial lawns. We purified one phage per plaque morphotype and combinations (host and date), leading to a final collection of over 1,200 phages. Phages were then re-isolated three times on the sensitive host to ensure purification. High-titer stocks (>10⁹ PFU/mL) were prepared via confluent lysis in agar overlays and stored at 4°C, with an additional aliquot stored at −80°C in the presence of 25% glycerol.

We prioritized sequencing phages targeting strains within clades V1 to V8, successfully obtaining over 1,000 new phage genomes (manuscript in preparation), none of which were identified as Schizotequatroviruses. To bridge this gap, we designed three primer sets (**Supplementary Table 1**) targeting conserved regions of Schizotequatrovirus genomes available in the NCBI RefSeq/GenBank database (**Supplementary Table 2**). These primers were tested on crude lysates of phages infecting strains outside clades V1–V8. This targeted approach enabled the detection of 17 phages, each yielding PCR products of the expected size with at least two primer pairs. Morphological analysis by electron microscopy and genome sequencing further confirmed their classification.

### Electron Microscopy

To characterize the morphology of the different phages, 20 μl of phage concentrates were adsorbed for 10 min to a formvar film on a carbon-coated 300 mesh copper grid (FF-300 Cu formvar square mesh Cu, delta microscopy). The adsorbed samples were negatively contrasted with 2% Uranyl acetate (EMS, Hatfield, PA, USA) before observation under a Jeol JEM-1400 Transmission Electron Microscope equipped with an Orious Gatan camera. Analyses were conducted by S. Le Panse at the platform MERIMAGE (Station Biologique, Roscoff, France).

### Phage genome sequencing and clustering

Phage suspensions (3 mL, >10⁹ PFU/mL) were concentrated using 1X PEG 8000 and 1M NaCl, incubated overnight at 4°C, and centrifuged at 4,500 rpm for 30 minutes at 4°C. The resulting pellet was resuspended in 300 µL SM buffer (100 mM NaCl, 8 mM MgSO₄·7H₂O, 50 mM Tris-Cl). Concentrated phages were then treated with DNAse (10 µL, 1000 units, Promega) and RNAse (2.5 µL, 3.5 mg/mL, Macherey-Nagel) at 37°C for 30 minutes. DNA extraction included protein lysis (0.02 M EDTA pH 8.0, 500 µg/mL proteinase K, 0.5% SDS) for 30 minutes at 55°C, phenol-chloroform extraction, and ethanol precipitation. DNA was visualized on a 0.7% agarose gel (50V, overnight at 4°C) and quantified using Qubit.

Phage sequencing was performed at the Biomics platform of the Pasteur Institute (Paris, France). DNA was fragmented using Covaris (target size: 500 bp), and libraries were prepared with the TruSeq™ DNA PCR-Free High Library Prep Kit. Due to inefficiency in adaptor ligation, an amplification step of 14 cycles was added using Illumina P7 and P5 primers (IDT). Sequencing was carried out on a MiSeq Micro v2 flow cell with paired-end 2×150 cycles. Reads were trimmed using Trimmomatic v0.39 (20) and assembled de novo with SPAdes v3.15.2. Contaminant contigs were filtered using the UniVec Database, and the resulting one-contig phage genome was manually linearized

Phages were clustered using VIRIDIC v1.0r3.6 with default settings (21). Intergenomic similarities were determined through BLASTN pairwise comparisons, with virus classification into family (≥50% similarity), genus (≥70% similarity), and species (≥95% similarity) ranks following ICTV genome identity thresholds.

### Annotation of phage genome

Genome annotation was performed by Pharokka version 1.7.2 (22) with Phanotate version 1.5.0 employed for coding sequence (CDS) identification (23). tRNAs genes were characterized using tRNAscan-SE 2.0 (24). Additional functional information was obtained through eggNOG-mapper version 2.1.12 (25), querying all unique proteins of the dataset. Putative defense systems were identified based on genes returned by the HMM search subprocess (26) of DefenseFinder 1.3.0 (27) in two independent runs: one sensitive (--no-cut-ga option) and the other specific (default parameters). The same strategy was applied to antidefense systems, using DefenseFinder with the --antidefensefinder-only option (28). Candidate receptor binding proteins were identified using PhageRBPdetection software version 3.0.0 (29) and one false positive among the eight hits was discarded due to an inconclusive AlphaFold-Multimer model (30). Finally, all genes from the reference genome 6E351A were manually curated and classified into the 14 functional categories presented in **Figure 2**.

**Figure 1.**
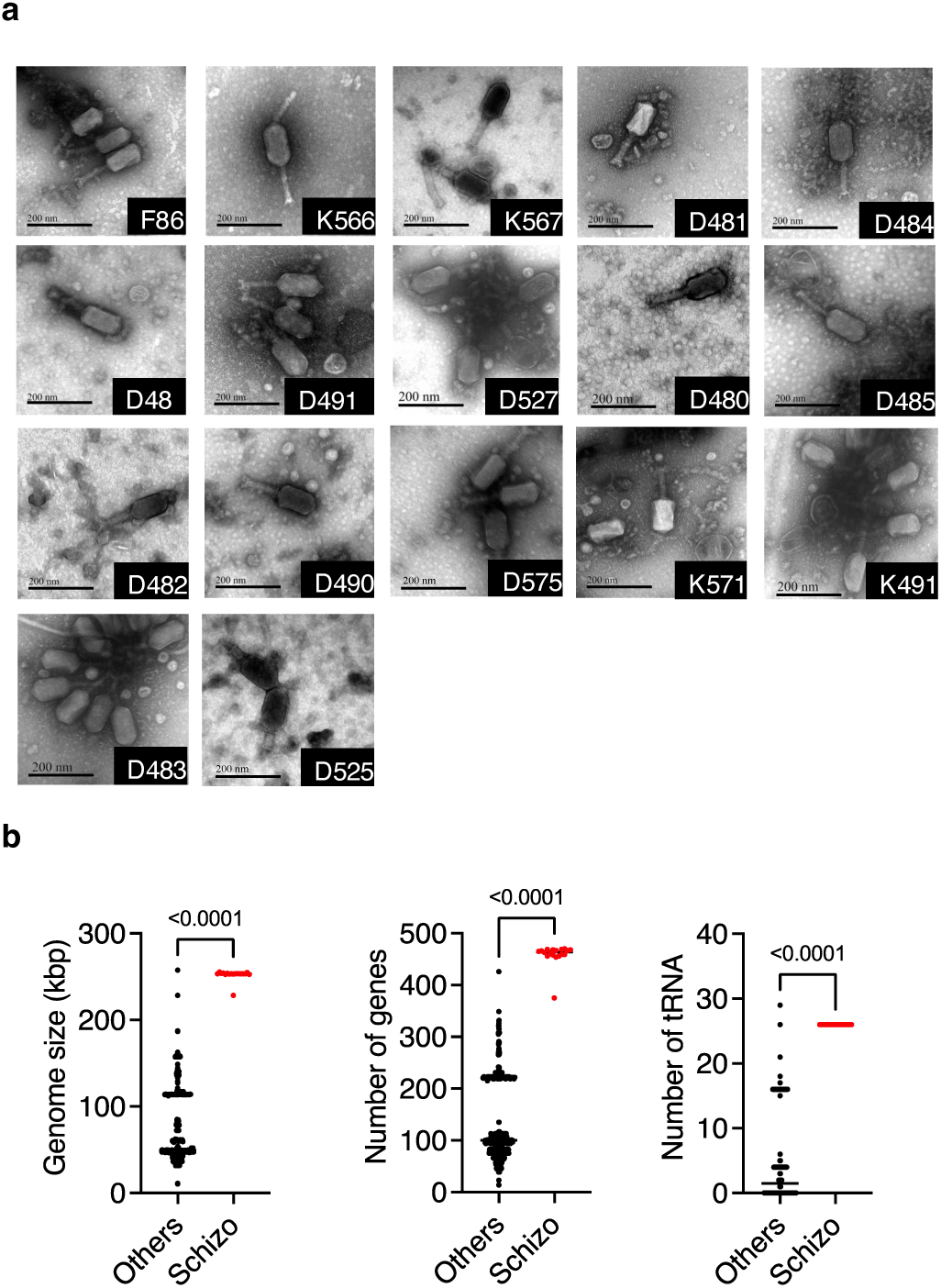
Morphological and genomic distinctions of newly isolated Schizotequatroviruses. **a** Transmission electron micrographs of 17 newly isolated Schizotequatroviruses produced using diverse *V. crassostreae* as host. Images are representative of the observation of dozens of particles. Images were collected using a Jeol JEM-1400 TEM Microscope. Scale bars (200 nm) are provided on each picture for reference. **b** Schizotequatroviruses exhibit distinctive characteristics including larger genome size, greater gene number, and numerous tRNAs. We compared genome size, predicted gene count, and tRNA presence between 18 Schizotequatroviruses (Schizo, represented by red dots) and the remaining 1,250 phages in our collection, all isolated from *Vibrio* hosts (others, represented by black dots). P-values from unpaired t-tests are provided.

**Figure 2.**
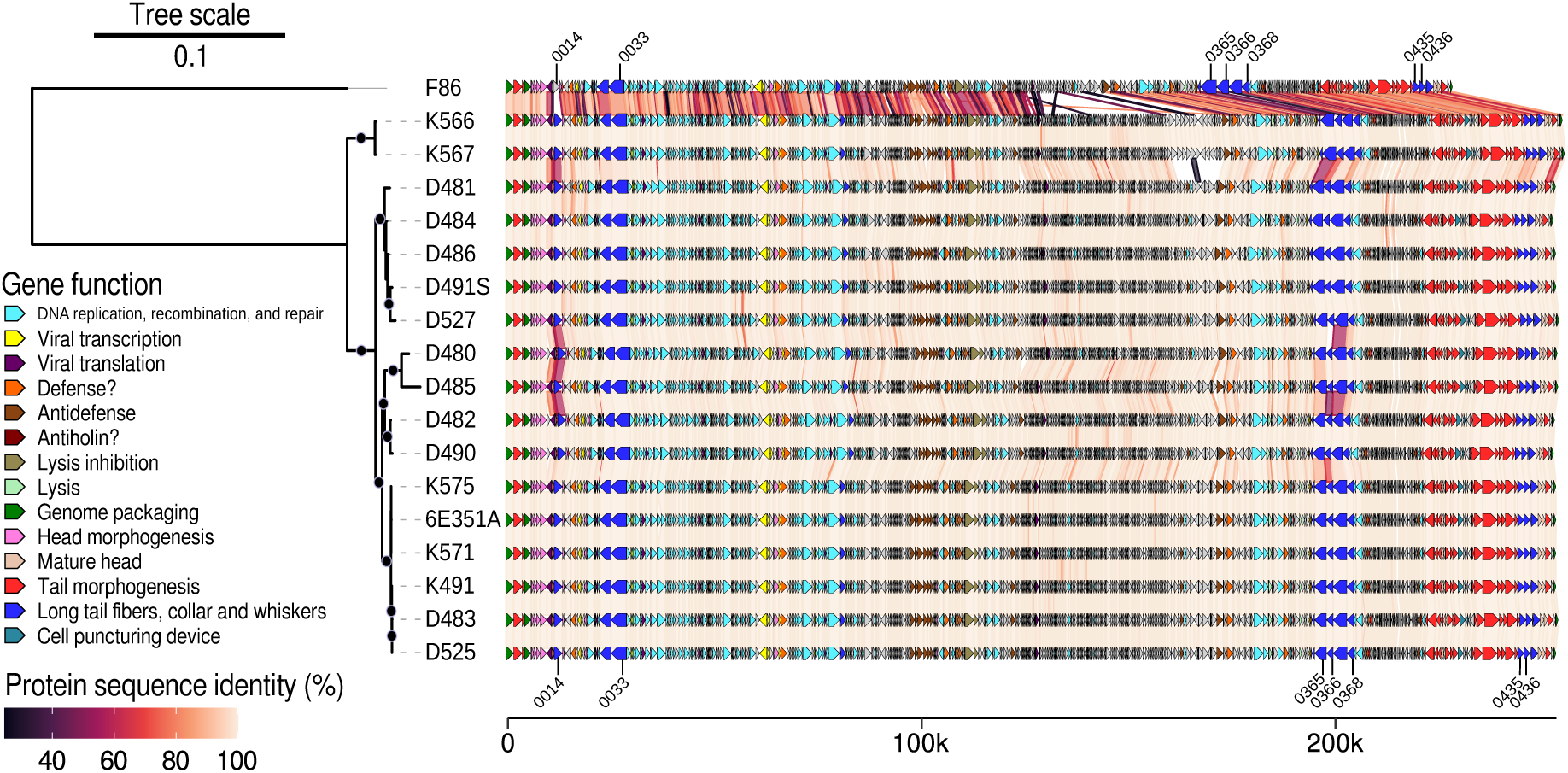
Phylogeny and synteny of Schizotequatroviruses. The maximum-likelihood core genome phylogeny of the phages is shown on the left, with branch lengths representing the estimated average number of amino-acid substitutions per site, as indicated by the scale bar at the top. Black dots indicate branches with over 90% bootstrap support. The visualization of the genomic organizations of phages were generated with the gggenomes R package with protein-coding genes colored according to the functional category of their inferred ortholog in the reference 6E351A genome (legend on the left). Genes of uncharacterized function are colored in grey. The scale at the bottom gives the genomic location of each gene (in base pairs). Synteny between pairs of phage genomes is represented by links connecting genes that encode proteins identified as bidirectional best hits (BBHs), with colors denoting their percentage of amino-acid sequence identity (legend in the bottom left corner). The seven groups of BBHs identified as coding for candidate receptor binding proteins (RBPs) are marked with a tick and labeled according to their position in phage 6E351A.

### tRNA analysis

The codon usage of Schizotequatroviruses and *V. crassostreae* was calculated based on the mean number of occurrences of each codon among the CDSs predicted for the 18 phages and the 157 host genomes, respectively. The anticodons and isotypes of phage tRNAs (notably the Met, fMet or Ile2 isotypes for the CAT anticodon) were extracted from the output GFF files generated by tRNAscan-SE 2.0. Anticodon-codon mapping was performed according to the pairing rules described in Table 1 from (31). The multipartite graph host codon / amino-acid / phage codon / phage anticodon was created using the sankeyNetwork function from the networkD3 R package version 0.4, with edge weights reflecting host codon usage, phage codon usage, and the mean copy number of phage tRNAs (1 copy of all tRNAs in all 18 phages except for tRNA_GTT_: 1 copy of tRNA_GTT_ in F86, 2 copies of tRNA_GTT_ in the 17 other phages).

### Core genome phylogenies

Maximum likelihood core genome phylogenies of *V. crassostreae* and Schizotequatroviruses were produced by PanACoTA version 1.4.1 (32), with different parameters to account for the differential level of genetic divergence between the two populations (*i.e.* species level for hosts with an average nucleotide identity >95% for any pair of genomes, family level according to VIRIDIC for phages). The host phylogeny was inferred at the DNA level by IQ-TREE (33), employing 1000 ultrafast bootstraps and the GTR+F+I+G4 model of nucleotide substitution, which was assessed as the best fit according to the Bayesian Information Criterion (BIC). This phylogeny was derived from the concatenated multiple sequence alignment (MSA) of 2,879 CDSs found in single copy in at least 99% of the genomes –this threshold was found to capture the majority of conserved single-copy gene families. Each marker set was defined as the CDSs of proteins sharing at least 80% amino-acid identity over at least 80% mutual length coverage, identified using the Linclust clustering algorithm from MMSeqs2 version 15-6f452 (34). The MSA of each protein cluster was generated using MAFFT version 7.525 (35) and converted into a CDS alignment via amino acid to codon mapping. The root was known from a previous work in which *V. gigantis* served as the outgroup (15). In contrast, the phylogeny of phages was inferred at the protein level (1000 bootstraps, LG+G4 as the best model of amino-acid substitution) from the concatenated MSA of 264 proteins present in single copy in at least 90% of the genomes. Each marker set was defined by Linclust as proteins sharing at least 25% sequence identity over at least 80% of their length. The tree was rooted with F86 as the outgroup, which was consistent with the root identified by both the midpoint and the minimal ancestor deviation unsupervised methods (36).

### Phages pangenome, synteny plot and fine-grained identification of core orthologs

Best-bidirectional hits (BBHs) were defined from an all-versus-all comparison of the 18 phage proteomes using MMseqs2 at maximal sensitivity (option -s 7.5), with an amino-acid identity cutoff of >20% and an alignment coverage of >50% of the length of the shortest protein. Groups of BBHs (BBH groups) were identified as connected components in the BBH graph, with 285 of the 547 BBH groups corresponding to core BBHs. 284 core BBH groups corresponded to single-copy BBHs while 1 core BBH group had a single copy in the F86 outgroup and 2 copies in the other 17 phages. For this group, we distinguished the two clusters of paralogs in the 17 phages using the OMA standalone orthology inference pipeline version 2.6.0 and only the paralogs found in the same OMA group as the unique protein of F86 were used in subsequent analyses of core genes. The pangenome graph was generated using the Python NetworkX version 3.2.1 package, with nodes representing BBH groups and edges indicating direct gene adjacency in at least one genome (weighted by the number of occurrences of the adjacency among the 18 genomes). The pangenome graph was visualized with Cytoscape version 3.10.0 (37) while the synteny plot was generated with the gggenomes R package version 0.9.12 (sequence identity scores of BBH pairs stored in the MMSeqs2 output were used to generate the links between proteins).

### Phylostratigraphy

To analyze events of gene loss, gain, and duplication, we leveraged the Hierarchical Orthologous Groups (HOGs) inferred during the previous run of the OMA standalone inference pipeline. As opposed to OMA groups which are inferred without a phylogeny and yields groups of orthologs with a unique member in each genome, a HOG is defined at each internal node of the recombination-free phylogeny and encompasses all descending paralogs and orthologs. HOGs are nested within HOGs of deeper levels in the phylogeny, up until the level of the “rootHOG”, namely the internal node of the phylogeny in which the gene family is inferred to have emerged / been gained. The 472 HOGs were mapped onto the nodes of the core genome recombination-free phylogeny using the ham function of pyHAM version 1.2.0 (Python HOGs Analysis Method) (38). The phylostratigraphy plot was generated with the create_tree_profile function of pyHAM.

### Recombination events and recombination-free phylogeny

For each group of core BBHs, proteins were aligned using MAFFT with the options –pairwise and –maxiterate 1000 for high accuracy. The corresponding codon alignments were generated through amino acid to codon mapping. The CDS alignments were then concatenated based on the gene order in the reference strain 6E351A. A preliminary core genome phylogeny was inferred from the concatenated CDS alignment using the GTR+G4 model of nucleotide substitution with 1000 bootstraps, and it was rooted with F86 as the outgroup. To identify recombination events and create a recombination-free phylogeny, Gubbins version 3.3.0 (39) was applied to the same alignment, using the aforementioned phylogeny as a starting tree and employing IQ-TREE with the GTR+G4 model of nucleotide substitution for tree construction. The topologies of recombination-unaware and -free phylogenies were compared using the Phylo.io online tool (40).

### Alignment-wide dN/dS

The MSA of proteins from each group of core orthologous genes was trimmed with Trimal version 1.4.1 to retain only sites with less than 20% gaps (41). The corresponding trimmed CDS alignments were generated using amino acid to codon mapping. For each trimmed CDS alignment, we applied Hyphy version 2.5.63 (42) to estimate a single alignment-wide dN/dS, using the recombination-free phylogeny as the guide tree and utilizing the recommended MG94CUSTOMCF3X4 model of codon substitution along with the GTR model of nucleotide substitution (command-line: `hyphy acd Universal $CDS_MSA MG94CUSTOMCF3X4 Global 012345 $RECOMBFREE_TREE Estimate`).

### Phages-bacteria interactions analyses

The heatmap in **Figure 3** was produced with the ComplexHeatmap R package version 2.15.4 (43). The specialization index of a phage is represented by its Paired Difference Index (PDI), as introduced by Poisot et al. in the context of bacteria-phage ecology (44). This index measures specificity as differential exploitation of bacterial isolates, by contrasting the highest infection score achieved on a host (PFU_1_) with those achieved on the other hosts (PFU_2_ to PFU_n_). The PDI for a given phage is therefore calculated as:

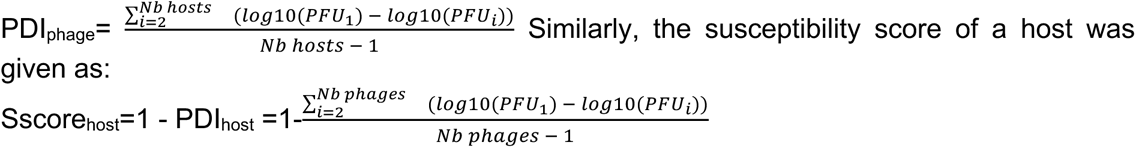

where PFU_1_ is the highest infection score recorded by a phage on the assessed host. Null PFU scores were treated as non-assigned values and PDI values were computed with the species level function of the R bipartite package. Hosts were classified into three distinct classes based on the susceptibility score distribution: resistant (score of 0), slightly susceptible (score above 0 but below 0.5), susceptible (score equal or higher than 0.5). The hypothesis that the distribution of either polymorphism (patristic distances), specificity to oysters, or specificity to the French West Coast of hosts with respect to their respective six closest strains differs across the three susceptibility classes was tested using the non-parametric Kruskal-Wallis test. Subsequent post-hoc pairwise tests were conducted using the non-parametric Mann-Whitney U test.

**Figure 3.**
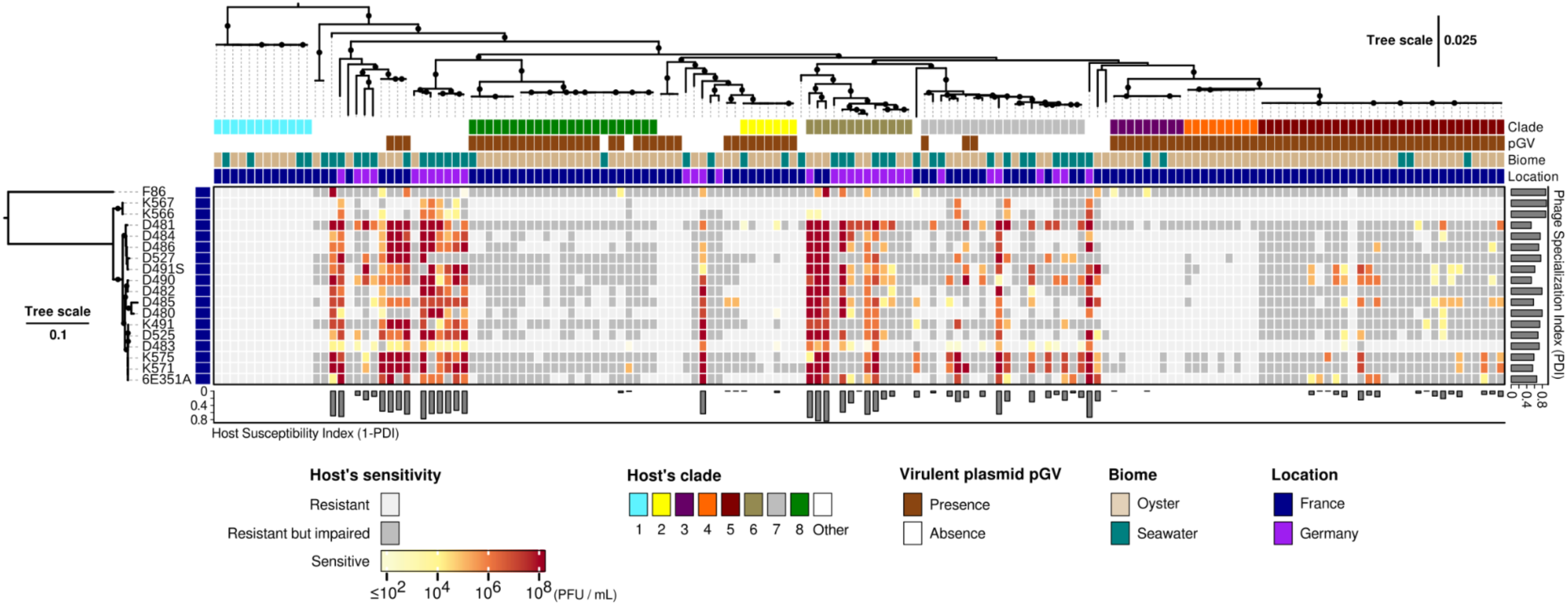
Schizotequatrovirus - *Vibrio crassostreae* quantitative interaction matrix. The central heat map displays the results of 2826 cross-infection assays (18 phages x 157 hosts). Rows represent phages, while columns represent hosts, ordered based on their respective core genome phylogeny. The scale of the phage phylogeny corresponds to the estimated average amino-acid substitution per site, while that of the host phylogeny corresponds to the estimated average nucleotide substitution per site. Four color strips aligned with the host phylogeny convey key information on the different *V. crassostreae* strains: 1) membership in the eight clades described in (15), 2) detection status of the virulent pGV plasmid (13), 3) environment of isolation (biome), and 4) sampling location. The color of each cell on the heatmap indicates the efficiency at which a phage can infect a host (in Plaques Forming Units (PFU) / mL), as indicated in the legend at the bottom left corner. A light grey cell signifies that the host is resistant to the phage, while a dark grey cell indicates that the host is impaired when challenged by high phage titers, but no production of viable phages is observed. The histogram on the right gives the specialization index of phages (PDI_phage_: Paired Difference Index), while that on the bottom gives the susceptibility index of hosts (1 – PDI_host_).

### Phylogenetic profiling of OmpK, LamB, GT and Wzy alleles

The OmpK, LamB, GT and Wzy families were identified as single linkage protein clusters (connected components in the sequence similarity graph) in the MMSeqs2 all-vs-all analysis of host proteomes, filtered for pairwise alignments yielding a least 25% identity and 70% mutual coverage. Identical proteins within each family were deduplicated at 100% sequence identity using the -c 1 option of CD-HIT version 4.8.1 (45). For each family, all unique protein variants were aligned with MAFFT, and a maximum-likelihood phylogeny was inferred from the MSA with IQ-TREE, utilizing the -m TEST option for finding the best model of amino-acid substitution (1000 bootstraps). Each tree was rooted using the midpoint method. Finally, for each protein family, a presence/absence matrix of variants across the host population was created. The matrix was visualized with the ComplexHeatmap R package (43) clustered according to the gene tree (rows) and the host phylogeny (columns).

### Structural analysis and prediction of the OmpK and LamB RBPs

To identify putative RBPs for OmpK and LamB, Alphafold-multimer (46) was utilized to perform pairwise structural predictions of complexes between OmpK and each of the seven candidate RBPs from phage 6E351A identified by PhageRBPdetection software (**Figure 2**), as well as LamB and the phage K566 orthologs of these seven candidate RBPs. The models of the OmpK-RBP complexes were unreliable, displaying non-physiologically relevant interaction interfaces and/or high inter-chain predicted aligned error (PAE) scores ≥ 30 Å, therefore no conclusions could be drawn from their analysis. However, several predicted LamB-RBP complexes were of sufficient quality (inter-chain PAE scores < 10 Å) that they could reflect a physiological receptor-RBP arrangement. To further narrow the list of candidates, the sequences of these RBPs were examined to identify K566/K567-specific variations that mapped to the predicted LamB-RBP interaction interface. Only one candidate, VPK566_0420/VPK567_0422, exhibited a difference in this region that could account for changes in receptor specificity, therefore this was assigned as the LamB RBP for phages K566 and K567.

Structures of *V. crassostreae* OmpK and LamB, as well as the seven candidate RBPs from phage 6E351A and K566, were predicted using AlphaFold2 (47) and AlphaFold-multimer as implemented in ColabFold (48), or AlphaFold3 (49) as appropriate. Identification of sequence-level conservation of protein domains was performed using CD-Search (50). Structural similarity searches against the Protein Data Bank (PDB) were performed using the Foldseek (51) and DALI (52) servers. Structural alignments were generated using the PDB Pairwise Structure Alignment tool (53). Calculation of electrostatic surface potential was performed using the Adaptive Poisson-Boltzmann Solver (APBS) service (54). Visualization of predicted structures was performed using ChimeraX (55). Multiple sequence alignments were generated using the default parameters of Muscle (version 5.1) as implemented in Geneious Prime 2025.0.2.

### Measurement of phage traits

Host range assays were carried out using an electronic multichannel pipette by spotting ten-fold serial dilutions onto host lawns. Plates were incubated overnight at RT and plaque formation was observed after 24 hours.

Phage adsorption experiments were conducted as described previously (15). Phages were mixed with exponentially growing cells (OD 0.3; 10⁷ CFU/mL) at an MOI of 0.01 and incubated at room temperature without agitation. At set intervals, 1mL of culture was transferred into a tube with 100 µL of chloroform, centrifuged at 14,000 rpm for 5 minutes. The supernatant was serially diluted and drop-spotted onto a sensitive host lawn to quantify remaining free phages.

Burst size and latency period were estimated using a protocol provided by the S. Moineau lab (Université Laval, Québec, Canada). Phages were mixed with exponentially growing cells (OD 0.3; 10⁷ CFU/mL) at an MOI of 0.01 and incubated at room temperature until maximum adsorption. Bacteria were then washed twice and the pellet was resuspended in a 1/100 dilution of fresh media. Every 10 minutes, an aliquot of the culture was collected and plated (either pure or diluted) onto a host lawn. After overnight incubation at room temperature, plaques were counted after 24 hours. Burst size was estimated as the average number of plaques at the first plateau divided by the average plaque count before the initial increase in plaque numbers (infection center). The latent period was defined as the time elapsed from initial phage adsorption to the first observed host cell lysis, determined as the last time point before PFU counts began to increase, plus the adsorption duration.

### Proteomic analysis of phage particles

Phage suspensions (10¹⁰ PFU) were precipitated using PEG as previously described, and the pellet was resuspended in 500 µL RIPA buffer (0.1% SDS, 1% Triton, 1 mM EDTA, 10 mM Tris-HCl, pH 8). Protein concentration was measured using the Biorad Protein Assay according to the manufacturer’s instructions, and samples were stored in 100 µL aliquots at −80°C. A total of 20 µg of protein was digested by trypsin and analyzed by tandem mass spectrometry. Peptides were mapped to the 6E351A phage proteome, and protein identities were assigned using the Protein Prophet algorithm (56) and validated with Scaffold (version Scaffold_5.3.3, Proteome Software Inc., Portland, OR). Peptide identifications were validated with a probability >80%, and protein identifications with a probability >90% with at least one identified peptide (57). Proteins with similar peptides that could not be differentiated were grouped. Data sources for protein quantification and identification are listed in **Supplementary Table 3**. Analyses were conducted by D. Faubert and M. Boulos, at the Proteomics Discovery Platform at the Montreal Clinical Research Institute, Canada.

### Detection of modified bases in phage DNA

DNA from phage 6E351A was purified using the Monarch® RNA Cleanup Kit (NEB), and 0.5 µg of DNA was hydrolyzed with a nucleoside digestion mix (NEB). The resulting nucleoside solution was analyzed by HPLC and mass spectrometry (LC/MS) (58). The HPLC-UV trace of 6E351A was aligned and compared to DNA digests from Bacillus phage PBP1 and phage Lambda. Analyses were conducted by Y.J. Lee in the P.R. Weigele lab, Biochemistry and Molecular Division, New England Biolabs, USA.

### Digital droplet PCR

A total of 2 mL of viral concentrates from seawater or plasma-derived viruses were used for DNA extraction (two sources, 35 dates, totaling 70 DNA samples). Phages were concentrated by adding 1X PEG 8000 and 1M NaCl, followed by overnight incubation at 4°C and centrifugation at 4,500 rpm for 30 minutes at 4°C. The resulting pellet was resuspended in 300 µL of SM buffer, DNA was then extracted following previously described protocols, resuspended in 100 µL and quantified using a Nanodrop, with an average concentration of 100 ng/µL.

Droplet digital PCR reactions consisted of 20 μL mixture per well containing 10 μL of ddPCR Evagreen Supermix, 600 nM of primers (**Supplementary Tables 1**) and 5 μL of DNA diluted 1/4. The ddPCR reactions were incorporated into droplets using the QX100 Droplet Generator (Bio-Rad). Nucleic acids were amplified with the following cycling conditions: 5 min at 95 °C, 40 cycles of 30 s at 95 °C and 60 °C for 60 s, and a final droplet cure step of 5 min at 4°C then 5 min at 90 °C using a Bio-Rad’s C1000 Touch Thermal Cycler. Droplets were read and analyzed using Bio-Rad QX600 system and QuantaSoft software (version 1.7.4.0917) in ‘absolute quantification’ mode. Only wells containing ≥10,000 droplets were accepted for further analysis.

### Molecular microbiology

Bacterial strains and plasmids used in this study are listed in **Supplementary Tables 4 and 5**. *V. crassostreae* isolates were grown on marine agar (MA) or broth (MB) at room temperature with gentle agitation, while *Escherichia coli* strains were cultured in Lysogeny Broth (LB) at 37°C with shaking. Where necessary, chloramphenicol (Cm; 5 or 25μg/mL for *V. crassostreae* and *E. coli*, respectively), thymidine (0.3 mM), and diaminopimelate (0.3 mM) were added to the media. The P_BAD_ promoter was induced with 0.2% L-arabinose and repressed with 1% D-glucose.

Cloning was performed with Gibson Assembly (New England Biolabs, NEB), with all constructs confirmed by sequencing. Gene inactivation was achieved using two methods: 1) Gene deletion was performed by cloning flanking regions into the pSW7848T suicide plasmid (59), allowing selection of mutants via P_BAD_-*ccdB* regulated integration and elimination steps, confirmed by PCR (60); 2) An internal region of the target gene was cloned into pSW23T (61), and single crossover integration was used for inactivation. For complementation, target genes or a *gfp* control were cloned under the P_LAC_ promoter in a P_MRB_ based vector (62) and introduced into mutants by conjugation.

Conjugation between *E. coli* and vibrios was carried out at 30°C on Tryptic soy agar (TSA) using a donor/recipient ratio of 5:1. ΔdapA donors were counter-selected on TSA without diaminopimelic acid but with antibiotics.

## RESULTS

### Isolation of Schizotequatroviruses

To expand our search for Schizotequatroviruses, we conducted a new time-series sampling at the same oyster farm (Brest, spanning 35 dates from July 1st to September 15th, 2021). This endeavor resulted in the generation of archives of virus concentrates from both seawater and oyster plasma. Utilizing our prior collection of *V. crassostreae* strains (15), we successfully isolated and sequenced 1044 viruses. We identified 17 new phages (1.7%) classified by the Virus Intergenomic Distance Calculator (VIRIDIC, (21)) within the Schizotequatrovirus family (**Supplementary Table 6**). Upon further examination through transmission electron microscopy (TEM), a striking morphological resemblance was observed between the 17 newly isolated Schizotequatroviruses (**Figure 1a**), 6E351A (15) and KVP40 (18). These phages were characterized by an icosahedral head approximately 125 nm in size and a contractile tail measuring about 110 nm. Genome sequencing unveiled an average genome length of 252,203 bp (+/- 5,983 bp) with 349–444 predicted coding sequences (CDSs) and precisely 26 tRNAs (**Figure 1b, Supplementary Table 7**).

### The large core genome of the Schizotequatrovirus

To characterize the variability of Schizotequatrovirus gene repertoires, we first identified putative orthologs as the genes that are bidirectional best hits (BBHs) between genomes. We then extracted the different connected components from the BBHs network to define groups of BBHs. On this basis, the pan-genome of the Schizotequatrovirus consisted of 543 gene families, of which 285 are families of core genes, *i.e.* with a member in all 18 genomes. Of relevance, most groups are either present in more than 80% of the genomes (76.6%) or in less than 20% (17.5%) (**Supplementary Figure 1**), comparable to what is observed in bacterial genomes (63). We then performed a computational characterization of gene functions in all genomes, followed by manual curation of the annotations in phage 6E351A. We could classify 178 BBH groups into 14 distinct functional categories (annotations of all BBH groups in **Supplementary Table 8**). A large number (48.6%) of core genes could not be classified because they have no known function.

One set of core genes encodes various components of viral particles (64), including the head, tail, long tail fibers, collar, and whiskers. A proteomic analysis of phage particles confirmed the function of the structural genes **(Supplementary Table 8)**. We identified no less than seven tail fibers (ranging from 474 to 1223 aa in length) using the PhageRBPdetection software that could be receptor binding proteins (RBPs) (29).

Another set of known persistent genes govern processes crucial for viral transcription, replication, procapsid assembly, genome packaging, tail assembly and host cell lysis to release the phage progeny **(Supplementary Table 8)**. Notably, and similarly to T4 phage, an RNA polymerase-ADP-ribosyltransferase, was detected in the viral particles. The T4 Alt gene product is a component of the phage head and enters the host cell during the process of infection together with the phage DNA to modulate T4 “early” promoter (65). Proteins involved in lysis (66) encompass an endolysin, a lytic transglycosylase, two putative holins and the inner and outer membrane spanin components. Lysis regulation could potentially involve the RIIA and B lysis inhibitors, two membrane-associated proteins shown to delay lysis through an as yet uncharacterized mechanism in T4 (67).

The core genome of the Schizotequatrovirus contains 26 tRNAs **(Figure 1b and Supplementary Table 8)**. The ability of a tRNA to decode a set of codons depends on post-transcriptional modifications which are hard to identify precisely from the genomes (68). Yet, assuming the right set of modifications, the 26 tRNAs could decode all 61 sense codons (**Supplementary Figure 2**). This set includes both fMet-tRNA and Met-tRNA, enabling the decoding of both the start and the standard ATG codons. The Schizotequatroviruses’ tRNAs are well aligned with the phage’s codon usage, since for 12 of the 18 amino-acids with synonymous codons, the most used codon can form a perfect Watson-Crick pair with the anticodon of a phage tRNA.

### The known repertoire of defenses and counter-defenses

The phage pan-genome includes numerous defense and counter-defense systems, aligning with the capacity of large-genome phages to accumulate such functions (**Supplementary Table 8**). The accessory genome of the Schizotequatrovirus includes 2 BBH groups of Viperins and a component of the PD-T4-9 defense system. Surprisingly, even though the repertoire of anti-defense systems usually evolves fast, we found several that are part of the core genome, e.g. the homologs of the NARP2 protein which circumvents NAD+ depletion of the Acb1 anti-CBASS protein and of the AcrIIA7 anti-CRISPR protein (28). We also identified a core 7-deazaguanine derivative biosynthesis gene cluster (69), which may play a role in restriction modification (R-M) system evasion. Four enzymes—FolE, QueD, QueE and QueC—are predicted to synthesize the precursor 7-cyano-7-deazaguanine (*preQ0*) from GTP. DpdA likely mediates the incorporation of *preQ0* into DNA as *dpreQ0*, a modified guanine base previously observed in Schizotequatrovirus phage nt-1 with 0.1% guanine modification (70). In phages containing QueF, an additional modification to 7-aminomethyl-7-deazaguanine (*preQ1*) occurs, modifying the DpdA substrate to preQ1. The presence of QueF in the 6E351A genome within this gene cluster suggests a dG modification to *dPreQ1* in its DNA. Mass spectrometric analysis of extracted DNA (**Supplementary Figure 3**) confirmed a 15% dG modification, with an observed mass of 295 Da, consistent with *dPreQ1*. Of note, we detected the FolE component of the 7-deazaguanine derivative biosynthesis gene cluster as well as the Acb1 Anti-CBASS and NARP2 NAD+ anti-depletion proteins in viral particles (**Supplementary Table 8**), hinting that the phage can deploy these antidefense proteins to immediately defend itself upon infection.

We examined the alignment between phage counter-defense genes and host defense systems in 21 host strains previously identified as susceptible to phage 6E351A (15). The 7-deazaguanine derivative biosynthesis gene cluster corresponded to the most frequently observed host defense systems, particularly R-M types I and II (**Supplementary Table 9**). Additionally, the Acb1, NARP2, and AcrIIA7 genes aligned with host defense systems such as CBASS_I, CBASS_II, and CAS Class 1-Subtype-I-F, which were each detected in at least one of the analyzed strains. With a total of 58 known defense systems identified across these susceptible strains, Schizotequatroviruses may harbor yet-undiscovered antidefense mechanisms capable of countering these diverse host defenses.

### The impact of recombination in Schizotequatroviruses

The previous results suggest a large conservation of gene composition among Schizotequatroviruses, with a few events of gene gains and losses. To better understand the evolution of these phages, we built a phylogenetic tree of a restricted set of single-copy gene families present in at least 17 of the 18 genomes, (**Figure 2**). The phylogeny of Schizotequatroviruses unambiguously flags phage F86 as the outgroup of the clade. This clade is further divided into two subclades: one composed of 15 phages, and the other containing the two phages K566 and K567. Having the phylogeny and the pangenome, we analyzed the conservation of genetic organization. The synteny appears remarkably conserved across the entire set of genomes, even when including the divergent F86 outgroup. Yet, the network of all gene adjacencies found in Schizotequatroviruses reveals a few highly variable regions in the pan-genome (forming loops in the network). These regions are enriched in uncharacterized proteins and in genes involved in defense/anti-defense systems, in line with their rapid turnover and their classification as accessory genes (**Supplementary Figure 1**).

When excluding the F86 outgroup, most BBHs appear nearly identical between phages (**Figure 2**). Yet, a substantial level of divergence is observed in a few loci (**Figure 2**). These loci correspond mostly to RBPs, which suggests that cell adsorption is a fast-evolving trait in these phages. We then wondered whether divergence within BBH groups resulted from homologous recombination events or rapid evolution by point mutations under diversifying selection. We first used Gubbins (39) to identify core genes with evidence of recombination, infer a recombination-free phylogeny, and then pinpoint the events of homologous recombination on each branch of the phylogenetic tree. Gubbins predicted 202 events of recombination on 13 internal and 16 terminal branches of the recombination-free phylogeny (**Supplementary Figure 4**). In total, 236 of the 285 loci present in all genomes were inferred to have undergone recombination during the diversification of the phage but some loci appeared to have recombined more than others (**Supplementary Figure 5**). The two main recombination hotspots in Schizotequatroviruses correspond to the 125-130kb and 195-205kb regions in 6E351A (**Figure 2**), which encode proteins of uncharacterized functions and long tail fiber RBPs, respectively. These two regions include core loci inferred to have undergone 9 recombination events during the diversification of the phage.

To assess if high sequence divergence is caused by episodes of positive selection, we estimated the gene-wide strength of selection acting against amino-acid substitution in core genes with Hyphy (42). This was obtained from the dN/dS ratio of nonsynonymous to synonymous substitution fixation rates. A dN/dS ratio below 1 suggests purifying selection, and a ratio higher than 1 suggests positive selection. All persistent genes had a dN/dS below 0.4 (**Supplementary Figure 5**). While this does not exclude positive selection at a few sites in the proteins, it does reveal a strong imprint of purifying selection inconsistent with the observation that some RBP BBHs share less than 60% identity in the clade of the 17 closely-related phages while the minimal pairwise identity in the vast majority of core BBH groups is above 90% (**Supplementary Table 8**). To confirm this conclusion, we compared the mean pairwise identity between BBHs within the most divergent core BBH groups (defined as a mean pairwise identity < 95% in the clade of 17 phages) and the number of inferred recombination events and the dN/dS ratio (**Supplementary Figure 6**). The results show a significant association of divergence with recombination and no association with the dN/dS ratio. This further suggests that divergence in core genes of Schizotequatroviruses is associated with recombination processes and not caused by selection for rapid accumulation of amino-acid substitutions from ancestral alleles.

In line with these findings, the branch lengths and topology of the phylogeny inferred at the DNA level from all persistent genes present in the 18 genomes is substantially different than the recombination-free phylogeny inferred by Gubbins (**Supplementary Figure 7**), and only the latter is congruent with the robust phylogeny inferred at the protein-level from conservative BBH groups (**Figure 2**). This affects the position of the phages K566 and K567 since recombination accounts for most nucleotide substitutions between their core genes and those of the 15 members of the sister subclade (**Supplementary Figure 7**). When recombination signals are removed by Gubbins, the two subclades appear to have diverged at the same rate of nucleotide substitution from their last common ancestor (**Supplementary Figure 7**). The inferred history of gene gains, duplications and losses (**Supplementary Figure 8**) identified another founding event of the shift of K566 and K567: the replacement of an ancestral anti-defense island (VP6E351A_0314 to _0326) by another one (VPK566_0302 to _0314 in phage K566) providing new functions, e.g. a glycosyltransferase and genes involved in Lipid A biosynthesis (VPK566_0309 and _0313 in **Supplementary Table 8**). Hence, the emergence of the sub-clade K566 and K567 is associated with events of recombination and acquisition of several genes. Together, these results suggest that recombination played a critical role in establishing polymorphism within Schizotequatroviruses.

### Variations in host range

To finely quantify the host range of the 18 Schizotequatroviruses isolated from *V. crassostreae*, we first measured the titer of each phage on 157 *V. crassostreae* strains, including 124 isolates from the same location (Brest, France) and 33 isolates from a different location (Sylt, Germany) (**Figure 3**). In sensitive hosts, the phage titers ranged from <10^2^ to 10^8^ PFU /mL. The host strain was classified as “resistant but impaired” if we observed a clearing zone but no production of viable phages and as “resistant” if no clearing zone nor plaque was observed.

We observed variability in host range and efficiency of infection across the Schizotequatroviruses population (**Figure 3**). We assessed the differential exploitation of bacterial isolates of each phage by computing the Paired Difference Index (PDI), a metric introduced to study specificity in the context of bacteria-phage ecology (44, 71). The higher a PDI is for a phage, the greater is the contrast between the highest infection score it can achieve on a host and the infection scores achieved on the other hosts. The F86 outgroup had a PDI of 0.89, K566/567 were found to be the most specialized with PDIs of 0.90 and 0.91, and the subclade of 15 phages had the broadest host range with an average PDI of 0.68 (**Figure 3**).

To broaden the diversity of population tested for host range, we compared the titer of phages 6E351A (representative of the subclade of 15 phages) and K567 (representative of the K566/567 subclade) on four additional *Vibrio* species (*V. cyclitrophicus, V. tasmaniensis, V. splendidus, and V. breoganii*) encompassing isolates from Brest or Plum Island (MA, USA) (72). Strains permitting phage reproduction were identified in all four species independently of their location (**Supplementary Figure 9a**). The phage pattern observed between strains of *Vibrio crassostreae* was reproduced for strains of the four other species, namely that K567 infected fewer strains of the same species than 6E351A (**Supplementary Figure 9b**).

We then analyzed the Schizotequatroviruses - *V. crassostreae* interaction matrix from the perspective of the infected host. We computed a susceptibility score for each host, given as 1 - PDI_host_, where the PDI of a host contrasts the highest infection score achieved by a phage on the host with the scores achieved by the other phages (**Figure 3**). The distribution of susceptibility scores clearly delineated three classes of hosts: resistant (score of 0), slightly susceptible (score above 0 but below 0.5) and susceptible (score of at least 0.5) (**Supplementary Figure 10a**). All strains from clade V1 were resistant to the 18 phages (**Figure 3**), suggesting that the diversification of this clade was accompanied by the loss of Schizotequatroviruses receptor(s). Strains from the ’patho-phylotypes’ (V2 to V5 and V8) carrying the virulence plasmid pGV (13, 16) were either resistant or slightly susceptible (**Figure 3**) although a good proportion of these strains were classified as resistant but impaired, indicating successful adsorption by the phages. The highest proportion of sensitive strains belonged to clades V6 and V7 or did not fall into any specific clades from (15) (**Figure 3**).

Intriguingly, the Schizotequatroviruses (all sampled in Brest) are more likely to infect clades of *V. crassostreae* which i) are characterized by a substantial polymorphism in core genes (**Supplementary Figure 10b**), ii) are not specific to Brest (**Supplementary Figure 10d**), and iii) are not found exclusively in oysters (**Supplementary Figure 10c**). Notably, strains from populations in locations without intensive oyster farming, such as Plum Island (USA) and Sylt (Germany), including *V. breoganii*, an algae specialist (73) were also susceptible to Schizotequatroviruses (**Supplementary Figure 9**). This provides an interesting ecological insight, namely that the generalist Schizotequatroviruses tend to target clades of generalist vibrios rather than clades of quasi-clonal patho-phylotype vibrios well adapted to a specific niche and known to be infected by specialist phages (15).

### OmpK and LamB are receptors for the Schizotequatroviruses

To better understand the patterns of *V. crassostreae* - Schizotequatroviruses interactions, we first sought to identify the attachment site of the phages on *V. crassostreae*. The attachment site can be a single type of receptor shared by diverse bacterial strains (monovalent phage) or correspond to multiple alternative receptors (polyvalent phage) (5).

The phage KVP40 receptor was previously identified as the outer membrane protein K (OmpK), a protein of unknown function constitutively expressed on the cell surface (74). Therefore, we investigated whether our collection of Schizotequatroviruses also requires OmpK to infect *V. crassostreae*. For 16 out of 18 phages the deletion of *ompK* in the host resulted in a resistant phenotype while the constitutive expression of *ompK* from a plasmid was sufficient to restore the infectivity of the phages (**Figure 4a**, **Supplementary Figure 11**). Adsorption assays further supported the hypothesis that OmpK is the receptor for phage 6E351A (**Figure 4b**). Consistent with these observations, the only clade of the *V. crassostreae* which is fully resistant to the Schizotequatrovirus (clade V1 in **Figure 3**) lacks the *ompK* gene (**Supplementary Figure 12**).

**Figure 4.**
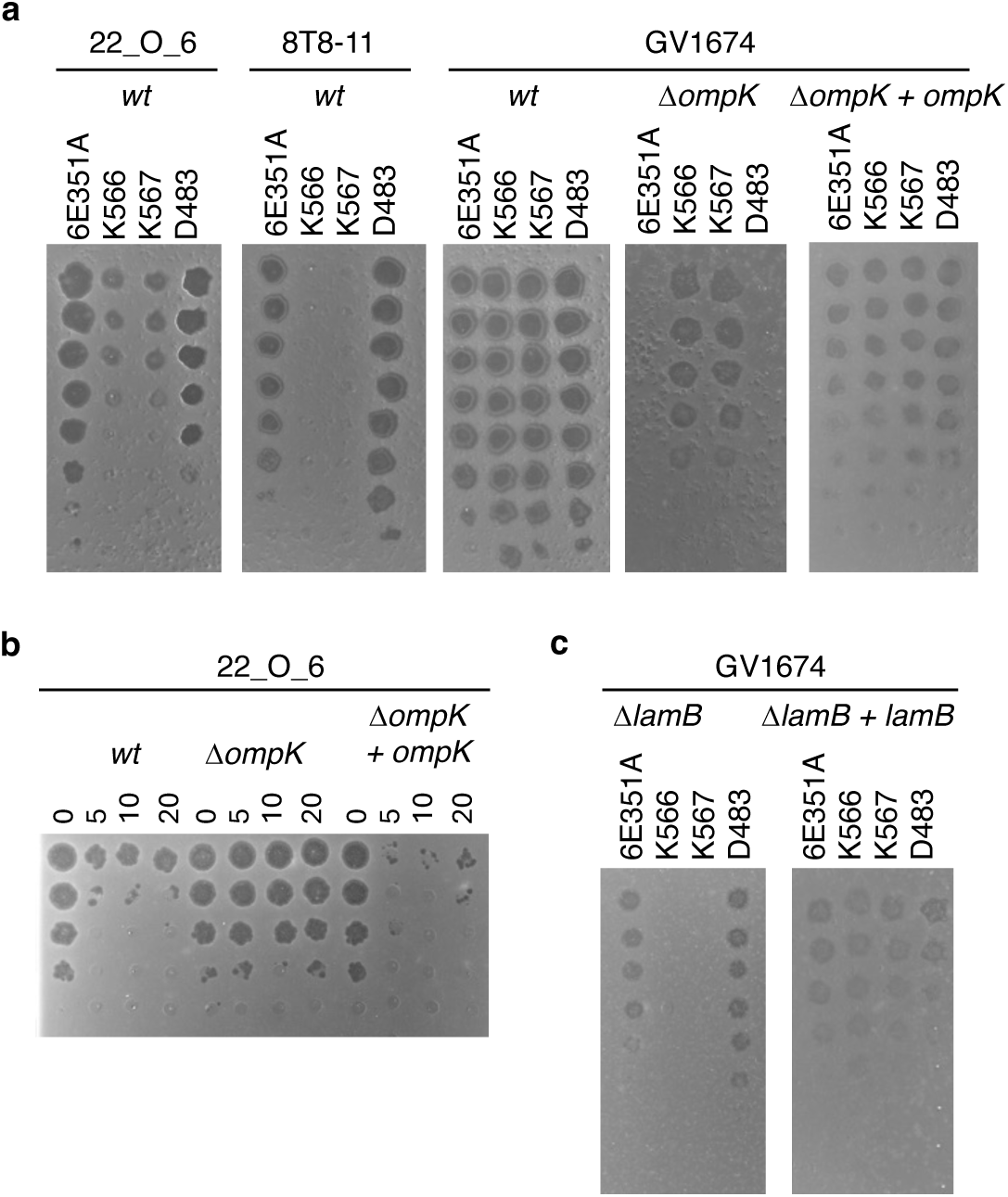
Role of OmpK and LamB as a receptor for Schizotequatroviruses. **a** Killing assays were performed by spotting 10-fold serial dilutions of each indicated phage onto the wild-type strain (wt), an *ΔompK* mutant derivative, and the *ΔompK* strain complemented with a plasmid constitutively expressing *ompK*. The tested strains are the original strains used for phage isolation: 22_O_6 for 6E351A, 8T8-11 for D483, and GV1674 for K566 and K567. **b** Adsorption assay for phage 6E351A on the wild-type strain 22_O_6 and its *ΔompK* mutant. A fixed concentration of phages (MOI 0.01) was allowed to adsorb to each strain for the indicated times, after which unattached (free) phages were serially diluted and plated with the original host, 22_O_6. A decrease in infectious particles indicates successful phage adsorption. **c** Killing assays performed on the *ΔlamB* mutant of GV1674 and its complemented derivative.

Yet, we found that phages K566 and K567 do not require OmpK for infection (**Figure 4a**). To identify their receptor(s), we generated spontaneous mutants from their original strain (GV1674) or its Δ*ompK* derivative. A total of 40 mutants were selected for genome sequencing, revealing 1 to 3 single nucleotide polymorphisms in the phage-resistant isolates (**Supplementary Table 10**). Among the 10 genes identified with nonsense mutations, frameshifts, or nonsynonymous mutations, two genes were notably affected, encoding the maltoporin LamB (10/40 strains) and its regulator MalT (20/40 strains). LamB has been previously reported as the main receptor for phages λ (75), T4-K10 (76), and T-even-like phage Tula (77). To confirm that LamB serves as a receptor for phages K566 and K567, we constructed a *lamB* deletion mutant in the original host strain (GV1674). The two phages failed to infect the single mutant Δ*lamB*, and the constitutive expression of *lamB* from a plasmid was sufficient to restore the infectivity of the phages (**Figure 4c**). We conclude than OmpK and LamB are receptors of this family of phages in *V. crassostreae*.

### Other host-encoded determinants underlie successful adsorption

After identifying OmpK and LamB as receptors for Schizotequatroviruses, we explored whether sequence diversity in these receptors across the host population could explain the varying infection success of each phage among the 157 *V. crassostreae* isolates. Phylogenetic profiling of OmpK and LamB variants within the host population (**Supplementary Figure 12**) revealed that the distribution of these variants does not account for the observed differences in phage adsorption profiles, as no receptor variant was specific to susceptible hosts. Instead, OmpK or LamB variants are found to be shared between both susceptible and resistant hosts. This is consistent with findings that OmpK phylogeny in *V. anguillarum* strains is not linked to susceptibility to phage KVP40 (78). Furthermore, when *ompK* or *lamB* were expressed in a fully resistant strain (31_O_70) from clade V1, the strain remained resistant to both phages 6E351A and K567 (**Figure 5a**). Together, these results indicate that while OmpK and LamB are necessary for attachment, they appear insufficient for infection by phages 6E351A and K567. Alternatively, the resistance of clade V1 strains could be explained by exopolysaccharides that effectively shield the cell surface, blocking phage adsorption (79).

**Figure 5.**
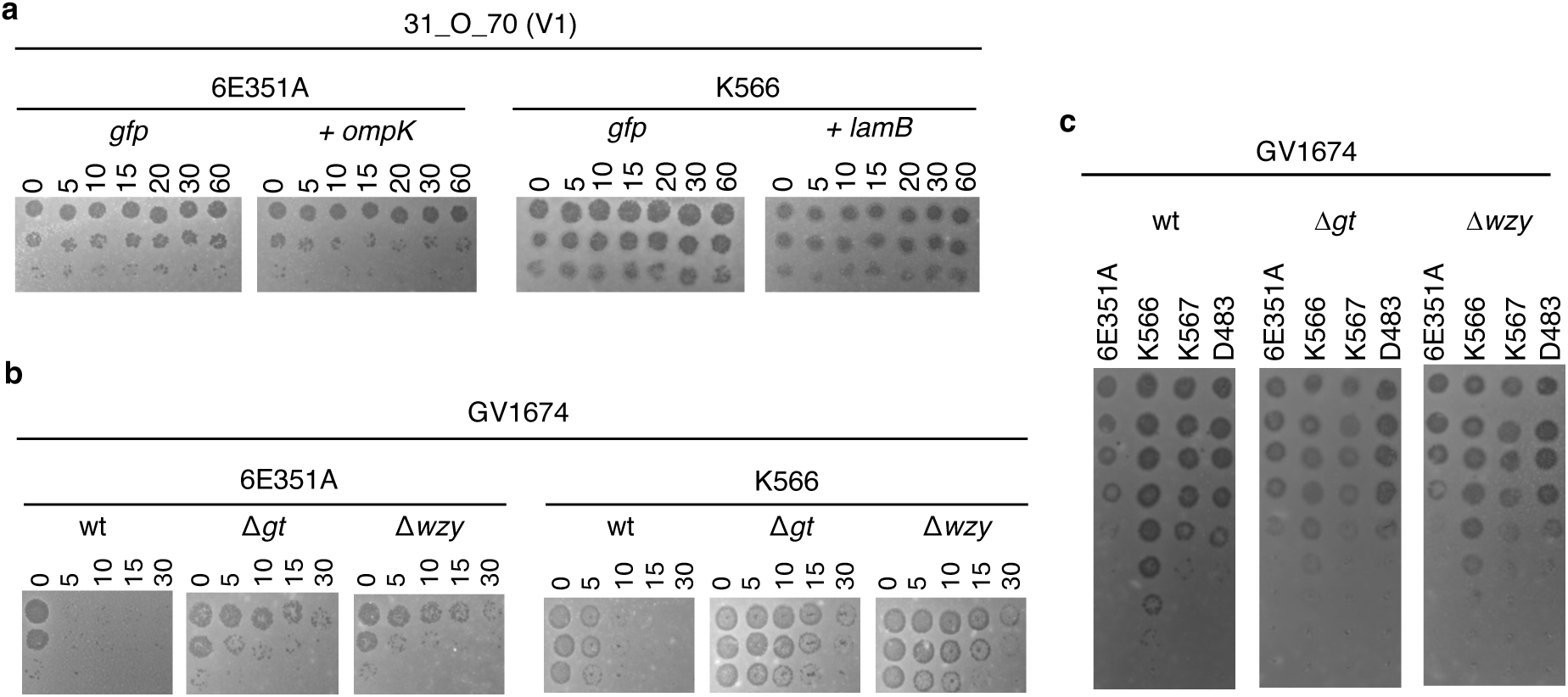
An EpsG family protein (Wzy) and a glycosyltransferase (GT) are involved in Schizotequatroviruses successful adsorption. **a** Killing assays were performed by spotting 10-fold serial dilutions of phage onto the strain 31_O_70 from clade V1 constitutively expressing the *gfp* (control), *ompK* or *lamB* from a plasmid. **b** Adsorption assay for phage 6E351A or K566 on the wild-type (wt) strain GV1674 or derivative in which we inactivated the *gt (τ<gt)* or *wzy (τ<wzy)* genes. A fixed concentration of phages (MOI 0.01) was allowed to adsorb to each strain for the indicated times, after which unattached (free) phages were serially diluted and plated with the original host, GV1674. A decrease in infectious particles indicates successful phage adsorption. **c** Killing assays were performed using strain GV1674 wt or derivative in which we inactivated the *gt* or *wzy* genes and 4 representative phages.

The infection cycle of most tailed phages begins with reversible attachment to an initial ’primary’ receptor on the host cell, often a sugar motif on surface glycans, followed by irreversible binding to a ’secondary’ receptor on the cell surface, which facilitates DNA injection (79). We hypothesized that, in addition to OmpK or LamB, an additional receptor might be involved in Schizotequatrovirus adsorption. Among the 40 phage-resistant isolates previously analyzed (**Supplementary Table 10**), two showed frameshifts in genes encoding a glycosyltransferase (*gt*, GV1674_v1_60037) and an EpsG family protein (*wzy*, GV1674_v1_60038), both located in a gene cluster involved in LPS synthesis. The *wzy* gene was unique to only 6 out of 157 *V. crassostreae* strains, while the glycosyltransferase gene was present in all strains with no allele specific to susceptible hosts (**Supplementary Figure 12**). Inactivation of *wzy* or *gt* was associated with a reduced adsorption rate for both phages 6E351A and K566 (**Figure 5b**). However, unlike inactivation of *ompK* or *lamB*, *wzy* or *gt* inactivation did not fully abolish infectivity (**Figure 5c**). This suggests that the LPS receptor may not be completely inactivated in these mutants or that these phages might utilize additional receptors, as has been demonstrated with phages from the Myoviridae morphotype (80). The presence of multiple receptors aligns well with our identification of seven loci encoding RBPs (**Figure 2**). This underscores the evolutionary flexibility of Schizotequatroviruses in expanding their host range.

### Candidate OmpK-, LamB- and LPS-binding proteins in Schizotequatroviruses

To gain insight into the similarities and differences between *V. crassostreae* OmpK, required for phage 6E351A recognition, and LamB, required for recognition by phages K566 and K567, we predicted their structures using AlphaFold2. OmpK was predicted to adopt a 12-stranded β-barrel (**Figure 6a** and **Supplementary Figure 13a**), with strong similarity to the monomeric *E. coli* nucleoside transporter Tsx (PDB 1TLW, rmsd 2.05 Å over 215 Cα). LamB was predicted to adopt a trimeric 18-stranded β-barrel (**Figure 6b** and **Supplementary Figure 13b**) with strong similarity to a *Salmonella typhimurium* maltoporin (PDB 1MPR, rmsd 2.4 Å over 348 Cα). We calculated the electrostatic surface potential of the extracellular-facing surfaces of OmpK and LamB, which revealed that LamB is significantly more electronegative than OmpK (**Figure 6c**). These electrostatic differences between the two receptors may correlate with complementary variations between the RBPs used by phages K566/K567 versus 6E351A.

**Figure 6.**
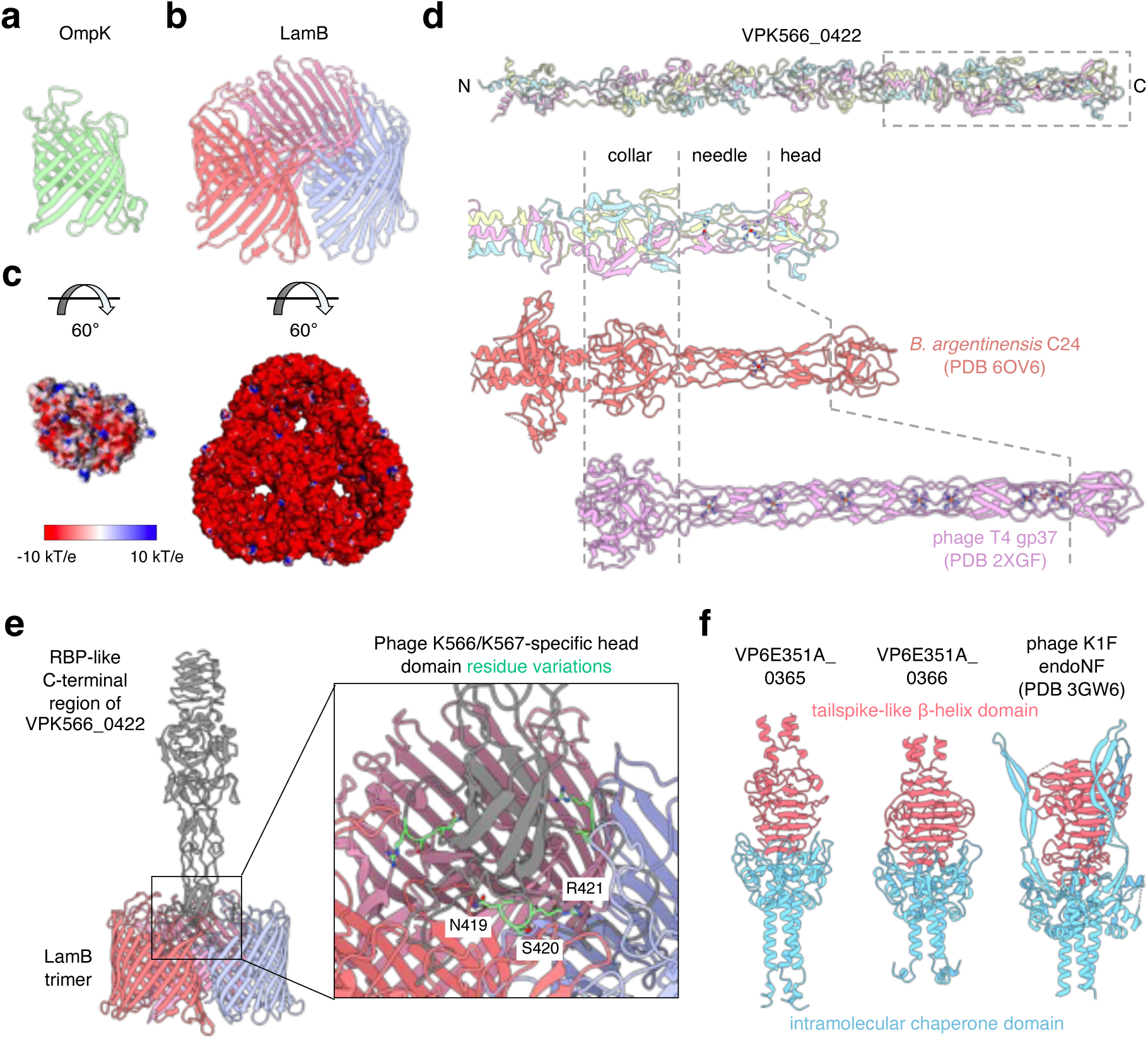
Phages K566 and K567 encode a putative RBP that may recognize LamB. **a** Structure of *V. crassostreae* OmpK predicted by AlphaFold2 (AF2). **b** Structure of *V. crassostreae* LamB predicted as a homotrimer by Alphafold-Multimer. **c** Electrostatic representation of the extracellular-facing surface of OmpK (left) and LamB (right) calculated using APBS; contoured from −10 kT/e (red) to 10 kT/e (blue). **d** (top) Structure of VPK566_0422 predicted as a homotrimer by AlphaFold3. The amino (N) and carboxy (C) termini are indicated. The grey dashed box indicates the receptor binding protein (RBP)-like region. (bottom) Comparison of the RBP-like region of VPK566_0422 to *Bizionia argentinensis* phage tip protein C24 (PDB 6OV6) and phage T4 long tail fiber tip protein gp37 (PDB 2XGF), drawn to scale. The shared collar, needle, and head domains of these proteins are indicated. Magnesium ions coordinated by the needle domains are depicted as red spheres **e** (left) Complex of LamB and VPK566_0422 (residues 295-476; grey), predicted by Alphafold-Multimer. (right) Close-up view of the head domain interaction with LamB. The phage K566- and K567-specific sequence variations are highlighted in green. **f** Comparison of the AlphaFold3-predicted trimeric structures of the C-terminal regions of VP6E351A_0365 and VP6E351A_0366 and the structure of the *E. coli* phage K1F endoNF tailspike (red) and intramolecular chaperone (blue) domains (PDB 3GW6).

To determine which RBP may be responsible for OmpK or LamB recognition, we examined the ability of the candidate RBPs to form a complex with OmpK or LamB using Alphafold-Multimer. While this approach was unable to identify a potential RBP for OmpK, VPK566_0422 (ortholog of VP6E351A_0436 from phage 6E351A) emerged as the strongest candidate for the LamB RBP in phages K566/K567. VPK566_0422 exhibits weak similarity to the receptor binding tip protein C24, belonging to a prophage in the genome of the Antarctic bacterium *Bizionia argentinensis* (81) (**Figure 6d, Supplementary Figure 14**). *B. argentinensis* C24 is itself structurally similar to the phage T4 long tail fiber receptor binding tip protein gp37, which recognizes the trimeric outer membrane β-barrel OmpC in *E. coli* (82). These three proteins share a comparable C-terminal architecture composed of a collar-, a magnesium coordinating needle-, and a head-like domain, albeit with significant variability in the length of the needle domains (**Figure 6d**). We identified sequence features unique to the K566 and K567 orthologs of VP6E351A_0436 in the region corresponding to the head domain (**Supplementary Figure 15a**), which were mapped onto the predicted structure of VPK566_0422 in complex with LamB (**Figure 6e, Supplementary Figure 15b**). These K566/K567-specific residues are optimally positioned to participate in bacterial receptor interactions, particularly the addition of an electropositive arginine residue which may be an adaptation to facilitate interactions with the more electronegative extracellular-facing surface of LamB (**Figure 6c**). The unique ability of phages K566 and K567 to use LamB for adsorption was probably the result of homologous recombination. Indeed, Gubbins inferred a single event of homologous recombination for the candidate LamB-RBP locus (VPK566_0422), precisely in the last common ancestor of K566 and K567 (**Supplementary Figure 4**).

To identify a potential RBP for LPS, we analyzed the sequences and predicted structures of the remaining RBPs (**Figure 2**). This revealed two candidates, VP6E351A_0365 and VP6E351A_0366, that exhibited sequence and structural similarity to the *E. coli* phage K1F tailspike endosialidase endoNF, which specifically binds to and degrades the polysialic acid capsule (83). Interestingly, VP6E351A_0365 and VP6E351A_0366 lack the sialidase domain, but have retained the tailspike-like triple β-helix as well as the C-terminal intramolecular chaperone domain that is required to fold the β-helix structure (**Figure 6f**, **Supplementary Figure 16a-b**). The tailspike domain of endoNF interacts with sialic acid-containing carbohydrates (83, 84), and similar β-helix structures in other phages are often linked to glycan interactions (85). Therefore VP6E351A_0365 and VP6E351A_0366 are strong candidates to fulfill the role of RBP for *V. crassostreae* LPS.

### Environmental abundance and life cycle dynamics of Schizotequatroviruses

The low frequency of Schizotequatrovirus in our collection of isolates (1.7%) is unexpected given their broad host range, and raises questions about the trade-offs between host range and other viral traits. Under culture conditions, broad host range phages with slower growth rates may be outcompeted by narrower host range phages with higher virulence (5). o address this potential bias, we employed a culture-independent technique, droplet digital PCR, to estimate the absolute abundance of phages in seawater and oyster plasma over the time series. When detected, Schizotequatrovirus genomes ranged from 57 to 470 copies per liter of seawater and 57 to 748 copies per milliliter of oyster plasma (**Supplementary Figure 17**). The abundance of Schizotequatroviruses was significantly lower than that of other phage genera, which specialize in phylo-pathotypes (**Figure 7a**). We conclude that despite their broad host range, extracellular Schizotequatroviruses’ relative abundance remains low in this environment.

**Figure 7.**
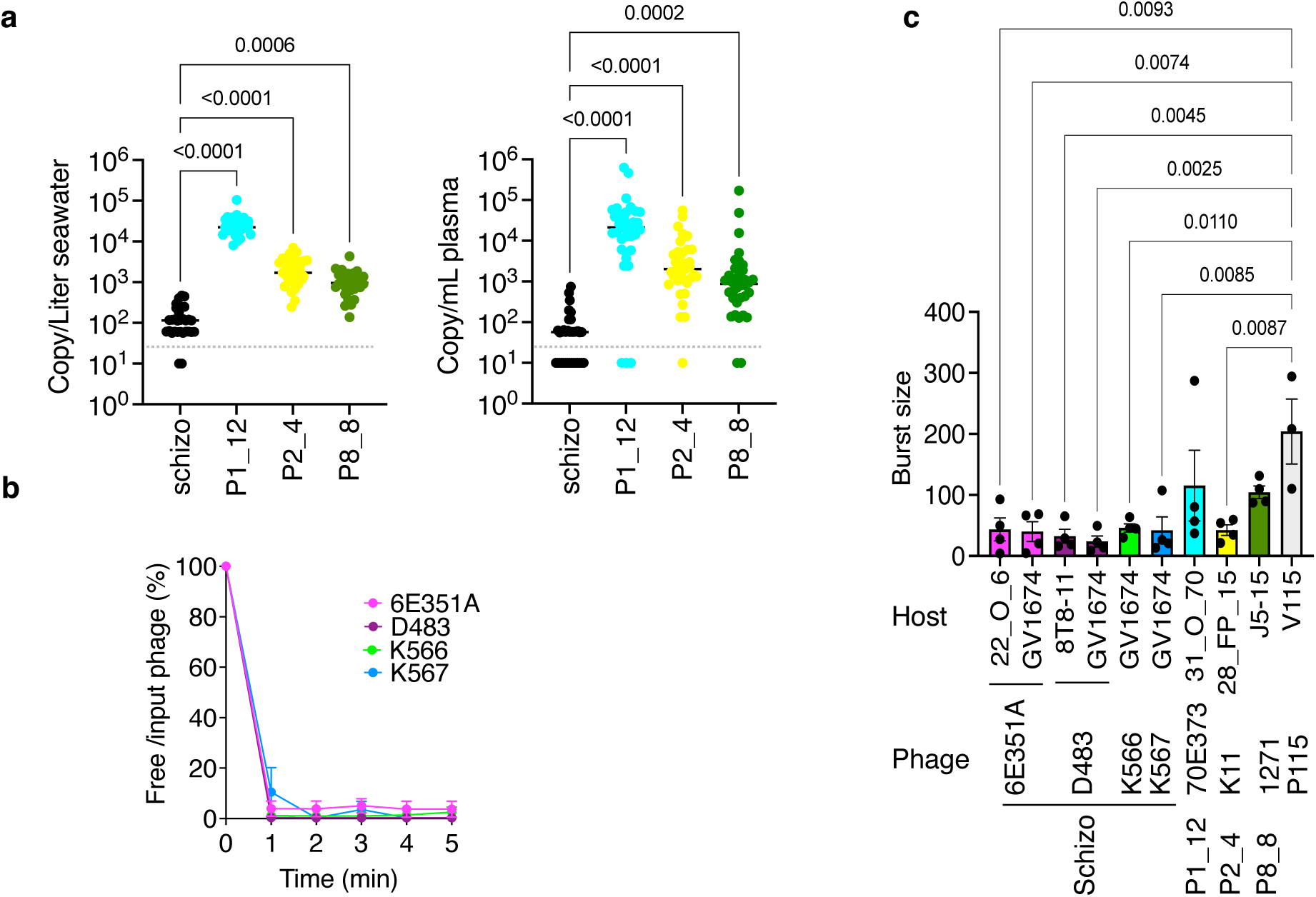
Environmental abundance and life cycle dynamics of Schizotequatroviruses. **a** Absolute abundance of Schizotequatroviruses and other phage genera (P1_12, P2_4, and P8_8, which infect *V. crassostreae* from clades V1, V2, and V8, respectively) in seawater and oyster plasma. Genome copies per Liter of seawater or mL of oyster plasma were quantified using droplet digital PCR (ddPCR). DNA samples were prepared from virome sources collected over 35 dates during the 2021 time series. The dashed line represents the detection limit. A Friedman test on RM one-way ANOVA results indicates that Schizotequatroviruses have significantly lower abundance compared to other phages in both oyster plasma and seawater. **b** Adsorption rates were calculated as the ratio of free phages to input phages at each time point. Data represent three independent experiments. **c** Burst sizes of the indicated phages were determined using one-step growth curves on original or alternative hosts (**Supplementary Figure 18**). Bars represent the mean ± standard error of the mean (SEM) from four independent experiments. Ordinary one-way ANOVA with Tukey’s multiple comparisons test shows a significant difference between all Schizotequatroviruses and P115, regardless of the host used to estimate burst size.

Host range shows a strong negative correlation with reproduction rate, while decay exhibits a positive correlation with reproduction rate, based on available data (86). The reproduction rate of a phage is primarily determined by three factors: adsorption rate (the speed at which a phage irreversibly binds to a host cell, initiating the infection process), latent period (the time from host cell infection to the release of new viruses), and burst size (the number of viral offspring produced per infected host cell). We therefore determined these parameters to identify which could be at a trade-off with host range and explain the low frequency of the Schizotequatrovirus.

Phage adsorption assays were performed by mixing phages with exponentially growing cells at a multiplicity of infection (MOI) of 0.01, followed by monitoring the loss of free phages over time. We previously observed that phage 6E351A rapidly adsorbed to its original host strain, 22_O_6, as well as to GV1674 (**Figures 4b and 5b**). This rapid adsorption rate (within 1 minute) was further confirmed with strain GV1674 using four representative Schizotequatroviruses (6E351A, D483, K566, and K567), regardless of whether their receptor was OmpK or LamB (**Figure 7b**).

The latent period and burst size were determined using a one-step growth curve with an MOI of 0.01. The latent period was estimated to be 30–40 minutes (**Supplementary Figure 18**), aligning with the bacterial generation time. The burst size averaged 38 ± 27 particles per cell, with no significant differences observed across phage-host combinations (**Figure 7c**, **Supplementary Figure 18**). Notably, the burst size of Schizotequatroviruses was similar to that of narrower-host-range phages from genera P1_12, P2_4, and P8_8, which infect *V. crassostreae* from clades V1, V2, and V8, respectively. However, phage P115, which is highly specific to a strain of *V. chagasii*, exhibited a significantly higher burst size (87, 88). Collectively, these findings indicate that the lower environmental abundance of Schizotequatroviruses compared to specialist phages targeting clades of *V. crassostreae* is not attributable to differences in burst size, as estimated *in vitro.* While these *in vitro* results fail to account for their relatively low abundance, they highlight the limitations of laboratory experiments in replicating the complexity of natural ecosystems. Future studies employing more “ecorealistic” approaches will be essential to uncovering the trade-offs between their broad host range and replication dynamics.

## DISCUSSION

Our study reveals the genetic mechanisms and evolutionary processes that enable Schizotequatroviruses to sustain a generalist lifestyle. These adaptations fall into two categories: persistent and flexible traits. Persistent traits include the ability to bind the widely distributed OmpK receptor within *Vibrionaceae*, as well as a viral RNA polymerase and tRNAs optimized for phage codon usage, which enhance transcriptional efficiency and host takeover. Additionally, a conserved 7-deazaguanine derivative biosynthesis gene cluster likely aids in evading diverse R-M systems.

The evolutionary flexibility of Schizotequatroviruses is exemplified by their acquisition and rapid turnover of multiple RBPs, enabling adaptation to receptor mutations, as well as antidefense systems. For instance, a lineage shift from the OmpK to LamB receptor, coupled with the incorporation of a novel antidefense island, demonstrates their capacity for adaptive evolution. Recombination emerges as a key mechanism supporting broad host range, particularly through the exchange of tail fibers, allowing phages to overcome selective pressures from host defenses and competition with more virulent specialists.

The relative scarcity of Schizotequatroviruses in the environment suggests they face fitness costs that offset the benefits of a broad host range (5, 89). *In vitro* experiments showed their rapid adsorption rates can be both an advantage and a liability—allowing quick host access but also risking attachment to non-viable targets, reducing fitness (86). While slow replication rates are observed, this trait is not unique to generalist phages, as some specialists also exhibit similar rates. Laboratory assessments, however, often overlook ecological factors like phage dispersion, host density, and spatial structure. Future studies should explore reproduction dynamics under ecologically relevant conditions and leverage single-cell analyses to unravel the sustainability paradox of these broad-host-range but environmentally rare phages.

We observed that Schizotequatroviruses tend to target clades of generalist hosts whose members are found among different geographic locations and biomes rather than the patho-phylotypes specifically adapted to oysters. These generalist vibrios, often avirulent in oysters, likely enter the oyster environment passively via filter-feeding and thrive in open seawater (13). Conversely, patho-phylotypes, which bloom in diseased oysters, are predominantly infected by specialist phages abundant in such niches (Manuscript in preparation). Oyster farming practices may actively select for narrow bacterial clades and their corresponding specialist phages, which are abundant in this environment. Investigating Schizotequatroviruses outside epidemic events may reveal a higher environmental prevalence and clarify their ecological role.

Schizotequatroviruses’ large genomes offer notable advantages, including the ability to encode diverse RBPs and antidefense mechanisms, enabling broad host range. These genomes also support regulatory complexity, optimizing infection based on host physiological states (90). However, large genomes might come with trade-offs: they require more resources and time for replication, resulting in slower replication rates (91–93), and may depend on sophisticated host machinery, limiting the range of potential hosts. Additionally, the increased genetic content presents more targets for host defenses like CRISPR-Cas systems. Exploring Schizotequatroviruses alongside other large-genome viruses could shed light on the interplay between genome size, host range, and evolutionary viability (94).

Considering long-term applications, such as preventing vibriosis in aquaculture, previous studies demonstrated the efficacy of KVP40 in reducing fish larval mortality (95), despite eventual resistance development (78, 96, 97). For juvenile oysters, our findings suggest that a single Schizotequatrovirus may not adequately cover the genetic diversity of *V. crassostreae* pathogens. A cocktail of specialist phages might provide more robust protection. Nonetheless, elucidating the molecular mechanisms underlying Schizotequatroviruses’ broad host range— particularly antidefense systems—holds promise for engineering phages tailored to specifically target phylo-pathotypes sustainably and effectively.

In conclusion, Schizotequatroviruses reveal complex trade-offs that may be inherent to generalist phages. Their broad host range, genomic adaptability, and ecological dynamics highlight both their potential and limitations in diverse bacterial landscapes. Combining insights from natural diversity with targeted engineering holds potential for developing biocontrol strategies that balance efficacy with sustainability, offering promising avenues for future applications in microbial management.

## ACKNOWLEDGEMENTS

We warmly thank Marc Monot and Laurence Ma (Biomics Platform, Institut Pasteur, France), Sophie Le Panse (MERIMAGE, Station Biologique de Roscoff, France), Peter Weigele and Yan-Jiun Lee (New England Biolabs, USA), and Denis Faubert and Marguerite Boulos (Proteomics Discovery Platform, Montreal Clinical Research Institute, Canada) for their invaluable technical assistance. We extend our gratitude to Bruno Petton (IFREMER, France) and other members of the GV team (Station Biologique de Roscoff, France) for their support during the time series sampling. Special thanks to Alice Perrault-Jolicoeur, Sylvain Moineau and his lab (Université Laval, Québec, Canada) for their insightful suggestions.

This work was supported by funding from the European Research Council (ERC) under the European Union’s Horizon 2020 research and innovation program (grant agreement No. 884988, Advanced ERC Dynamic), the Canada Excellence Research Chairs Program (CERC-2022-00051), and the Fonds de recherche du Québec – Nature et technologies (FRQ-NT, Fonds des leaders John-R.-Evans, grant 44584) awarded to FLR. Substantial support was provided by the Agence Nationale de la Recherche (ANR-20-CE35-0014 “RESISTE”) to EPCR and FLR, as well as by INCEPTION PIA/ANR-16-CONV-0005 and Laboratoire d’Excellence IBEID (Integrative Biology of Emerging Infectious Diseases, ANR-10-LABX-62-IBEID) for EPCR. PD was supported by the ANR (ANR-19-CE20-0004 “DECICOMP”). YVB was supported by a Canada 150 Research Chair.

This work utilized the computational and storage services (TARS cluster) provided by the IT department at Institut Pasteur, Paris.

## COMPETINGS INTERESTS

Authors declare no competing interests.

## DATA AVAILABILITY STATEMENT

### Data availability

The sequenced Schizotequatrovirus genomes have been deposited in the NCBI database, with accession numbers listed in Supplementary Table 7.

### Materials availability

All vibrio strains, phage strains and plasmids are available upon request.

## REFERENCES

1. Breitbart M, Bonnain C, Malki K, Sawaya NA. Phage puppet masters of the marine microbial realm. Nat Microbiol. 2018;3(7):754–66.

2. Ofir G, Sorek R. Contemporary Phage Biology: From Classic Models to New Insights. Cell. 2018;172(6):1260–70.

3. Salmond GP, Fineran PC. A century of the phage: past, present and future. Nat Rev Microbiol. 2015;13(12):777–86.

4. Strathdee SA, Hatfull GF, Mutalik VK, Schooley RT. Phage therapy: From biological mechanisms to future directions. Cell. 2023;186(1):17–31.

5. de Jonge PA, Nobrega FL, Brouns SJJ, Dutilh BE. Molecular and Evolutionary Determinants of Bacteriophage Host Range. Trends Microbiol. 2019;27(1):51–63.

6. Holtappels D, Alfenas-Zerbini P, Koskella B. Drivers and consequences of bacteriophage host range. FEMS Microbiol Rev. 2023;47(4).

7. Georjon H, Bernheim A. The highly diverse antiphage defence systems of bacteria. Nat Rev Microbiol. 2023;21(10):686–700.

8. Koskella B, Hernandez CA, Wheatley RM. Understanding the Impacts of Bacteriophage Viruses: From Laboratory Evolution to Natural Ecosystems. Annu Rev Virol. 2022;9(1):57–78.

9. Dekel-Bird NP, Sabehi G, Mosevitzky B, Lindell D. Host-dependent differences in abundance, composition and host range of cyanophages from the Red Sea. Environ Microbiol. 2015;17(4):1286–99.

10. Kauffman KM, Hussain FA, Yang J, Arevalo P, Brown JM, Chang WK, et al. A major lineage of non-tailed dsDNA viruses as unrecognized killers of marine bacteria. Nature. 2018;554(7690):118–22.

11. Munson-McGee JH, Peng S, Dewerff S, Stepanauskas R, Whitaker RJ, Weitz JS, et al. A virus or more in (nearly) every cell: ubiquitous networks of virus-host interactions in extreme environments. ISME J. 2018;12(7):1706–14.

12. de Lorgeril J, Lucasson A, Petton B, Toulza E, Montagnani C, Clerissi C, et al. Immune-suppression by OsHV-1 viral infection causes fatal bacteraemia in Pacific oysters. Nat Commun. 2018;9(1):4215.

13. Bruto M, James A, Petton B, Labreuche Y, Chenivesse S, Alunno-Bruscia M, et al. Vibrio crassostreae, a benign oyster colonizer turned into a pathogen after plasmid acquisition. ISME J. 2017;11(4):1043–52.

14. Lemire A, Goudenège D, Versigny T, Petton B, Calteau A, Labreuche Y, et al. Populations, not clones, are the unit of vibrio pathogenesis in naturally infected oysters. The ISME journal. 2015;9(7):1523–31.

15. Piel D, Bruto M, Labreuche Y, Blanquart F, Goudenege D, Barcia-Cruz R, et al. Phage-host coevolution in natural populations. Nat Microbiol. 2022.

16. Piel D, Bruto M, James A, Labreuche Y, Lambert C, Janicot A, et al. Selection of Vibrio crassostreae relies on a plasmid expressing a type 6 secretion system cytotoxic for host immune cells. Environ Microbiol. 2020;22(10):4198–211.

17. Miller ES, Heidelberg JF, Eisen JA, Nelson WC, Durkin AS, Ciecko A, et al. Complete genome sequence of the broad-host-range vibriophage KVP40: comparative genomics of a T4-related bacteriophage. J Bacteriol. 2003;185(17):5220–33.

18. Matsuzaki S, Tanaka S, Koga T, Kawata T. A broad-host-range vibriophage, KVP40, isolated from sea water. Microbiol Immunol. 1992;36(1):93–7.

19. Kauffman KM, Polz MF. Streamlining standard bacteriophage methods for higher throughput. MethodsX. 2018;5:159–72.

20. Bolger AM, Lohse M, Usadel B. Trimmomatic: a flexible trimmer for Illumina sequence data. Bioinformatics. 2014;30(15):2114–20.

21. Moraru C, Varsani A, Kropinski AM. VIRIDIC-A Novel Tool to Calculate the Intergenomic Similarities of Prokaryote-Infecting Viruses. Viruses. 2020;12(11).

22. Bouras G, Nepal R, Houtak G, Psaltis AJ, Wormald PJ, Vreugde S. Pharokka: a fast scalable bacteriophage annotation tool. Bioinformatics. 2023;39(1).

23. McNair K, Zhou C, Dinsdale EA, Souza B, Edwards RA. PHANOTATE: a novel approach to gene identification in phage genomes. Bioinformatics. 2019;35(22):4537–42.

24. Chan PP, Lin BY, Mak AJ, Lowe TM. tRNAscan-SE 2.0: improved detection and functional classification of transfer RNA genes. Nucleic Acids Res. 2021;49(16):9077–96.

25. Cantalapiedra CP, Hernandez-Plaza A, Letunic I, Bork P, Huerta-Cepas J. eggNOG-mapper v2: Functional Annotation, Orthology Assignments, and Domain Prediction at the Metagenomic Scale. Mol Biol Evol. 2021;38(12):5825–9.

26. Eddy SR. Accelerated Profile HMM Searches. PLoS Comput Biol. 2011;7(10):e1002195.

27. Tesson F, Herve A, Mordret E, Touchon M, d’Humieres C, Cury J, et al. Systematic and quantitative view of the antiviral arsenal of prokaryotes. Nat Commun. 2022;13(1):2561.

28. Tesson F, Huiting E, Wei L, Ren J, Johnson M, Planel R, et al. Exploring the diversity of anti-defense systems across prokaryotes, phages, and mobile genetic elements. bioRxiv. 2024:2024.08.21.608784.

29. Boeckaerts D, Stock M, De Baets B, Briers Y. Identification of Phage Receptor-Binding Protein Sequences with Hidden Markov Models and an Extreme Gradient Boosting Classifier. Viruses. 2022;14(6).

30. Evans R, O’Neill M, Pritzel A, Antropova N, Senior A, Green T, et al. Protein complex prediction with AlphaFold-Multimer. bioRxiv. 2022:2021.10.04.463034.

31. Murphy FVt, Ramakrishnan V. Structure of a purine-purine wobble base pair in the decoding center of the ribosome. Nat Struct Mol Biol. 2004;11(12):1251–2.

32. Perrin A, Rocha EPC. PanACoTA: a modular tool for massive microbial comparative genomics. NAR Genom Bioinform. 2021;3(1):lqaa106.

33. Minh BQ, Schmidt HA, Chernomor O, Schrempf D, Woodhams MD, von Haeseler A, et al. IQ-TREE 2: New Models and Efficient Methods for Phylogenetic Inference in the Genomic Era. Mol Biol Evol. 2020;37(5):1530–4.

34. Steinegger M, Soding J. Clustering huge protein sequence sets in linear time. Nat Commun. 2018;9(1):2542.

35. Katoh K, Standley DM. MAFFT multiple sequence alignment software version 7: improvements in performance and usability. Mol Biol Evol. 2013;30(4):772–80.

36. Tria FDK, Landan G, Dagan T. Phylogenetic rooting using minimal ancestor deviation. Nat Ecol Evol. 2017;1:193.

37. Shannon P, Markiel A, Ozier O, Baliga NS, Wang JT, Ramage D, et al. Cytoscape: a software environment for integrated models of biomolecular interaction networks. Genome Res. 2003;13(11):2498–504.

38. Train CM, Pignatelli M, Altenhoff A, Dessimoz C. iHam and pyHam: visualizing and processing hierarchical orthologous groups. Bioinformatics. 2019;35(14):2504–6.

39. Croucher NJ, Page AJ, Connor TR, Delaney AJ, Keane JA, Bentley SD, et al. Rapid phylogenetic analysis of large samples of recombinant bacterial whole genome sequences using Gubbins. Nucleic Acids Res. 2015;43(3):e15.

40. Robinson O, Dylus D, Dessimoz C. Phylo.io: Interactive Viewing and Comparison of Large Phylogenetic Trees on the Web. Mol Biol Evol. 2016;33(8):2163–6.

41. Capella-Gutierrez S, Silla-Martinez JM, Gabaldon T. trimAl: a tool for automated alignment trimming in large-scale phylogenetic analyses. Bioinformatics. 2009;25(15):1972–3.

42. Kosakovsky Pond SL, Poon AFY, Velazquez R, Weaver S, Hepler NL, Murrell B, et al. HyPhy 2.5-A Customizable Platform for Evolutionary Hypothesis Testing Using Phylogenies. Mol Biol Evol. 2020;37(1):295–9.

43. Gu Z. Complex heatmap visualization. Imeta. 2022;1(3):e43.

44. Poisot T, Lepennetier G, Martinez E, Ramsayer J, Hochberg ME. Resource availability affects the structure of a natural bacteria-bacteriophage community. Biol Lett. 2011;7(2):201–4.

45. Li W, Godzik A. Cd-hit: a fast program for clustering and comparing large sets of protein or nucleotide sequences. Bioinformatics. 2006;22(13):1658–9.

46. Evans R, O’Neill M, Pritzel A, Antropova N, Senior A, Green T, et al. Protein complex prediction with AlphaFold-Multimer. bioRxiv. 2021:2021.10.04.463034.

47. Jumper J, Evans R, Pritzel A, Green T, Figurnov M, Ronneberger O, et al. Highly accurate protein structure prediction with AlphaFold. Nature. 2021;596(7873):583–9.

48. Mirdita M, Schutze K, Moriwaki Y, Heo L, Ovchinnikov S, Steinegger M. ColabFold: making protein folding accessible to all. Nat Methods. 2022;19(6):679–82.

49. Abramson J, Adler J, Dunger J, Evans R, Green T, Pritzel A, et al. Accurate structure prediction of biomolecular interactions with AlphaFold 3. Nature. 2024;630(8016):493–500.

50. Chen Y, Nie F, Xie SQ, Zheng YF, Dai Q, Bray T, et al. Efficient assembly of nanopore reads via highly accurate and intact error correction. Nat Commun. 2021;12(1):60.

51. van Kempen M, Kim SS, Tumescheit C, Mirdita M, Lee J, Gilchrist CLM, et al. Fast and accurate protein structure search with Foldseek. Nat Biotechnol. 2024;42(2):243–6.

52. Holm L, Laiho A, Toronen P, Salgado M. DALI shines a light on remote homologs: One hundred discoveries. Protein Sci. 2023;32(1):e4519.

53. Bittrich S, Segura J, Duarte JM, Burley SK, Rose Y. RCSB protein Data Bank: exploring protein 3D similarities via comprehensive structural alignments. Bioinformatics. 2024;40(6).

54. Jurrus E, Engel D, Star K, Monson K, Brandi J, Felberg LE, et al. Improvements to the APBS biomolecular solvation software suite. Protein Sci. 2018;27(1):112–28.

55. Meng EC, Goddard TD, Pettersen EF, Couch GS, Pearson ZJ, Morris JH, et al. UCSF ChimeraX: Tools for structure building and analysis. Protein Sci. 2023;32(11):e4792.

56. Nesvizhskii AI, Keller A, Kolker E, Aebersold R. A statistical model for identifying proteins by tandem mass spectrometry. Anal Chem. 2003;75(17):4646–58.

57. Keller A, Nesvizhskii AI, Kolker E, Aebersold R. Empirical statistical model to estimate the accuracy of peptide identifications made by MS/MS and database search. Anal Chem. 2002;74(20):5383–92.

58. Lee YJ, Weigele PR. Detection of Modified Bases in Bacteriophage Genomic DNA. Methods Mol Biol. 2021;2198:53–66.

59. Val ME, Skovgaard O, Ducos-Galand M, Bland MJ, Mazel D. Genome engineering in Vibrio cholerae: a feasible approach to address biological issues. PLoS genetics. 2012;8(1):e1002472.

60. Le Roux F, Binesse J, Saulnier D, Mazel D. Construction of a Vibrio splendidus mutant lacking the metalloprotease gene vsm by use of a novel counterselectable suicide vector. Appl Environ Microbiol. 2007;73(3):777–84.

61. Demarre G, Guerout AM, Matsumoto-Mashimo C, Rowe-Magnus DA, Marliere P, Mazel D. A new family of mobilizable suicide plasmids based on broad host range R388 plasmid (IncW) and RP4 plasmid (IncPalpha) conjugative machineries and their cognate Escherichia coli host strains. Res Microbiol. 2005;156(2):245–55.

62. Le Roux F, Davis BM, Waldor MK. Conserved small RNAs govern replication and incompatibility of a diverse new plasmid family from marine bacteria. Nucleic Acids Res. 2011;39(3):1004–13.

63. Francis AR, Tanaka MM. Evolution of variation in presence and absence of genes in bacterial pathways. BMC Evol Biol. 2012;12:55.

64. Yap ML, Rossmann MG. Structure and function of bacteriophage T4. Future Microbiol. 2014;9(12):1319–27.

65. Koch T, Raudonikiene A, Wilkens K, Ruger W. Overexpression, purification, and characterization of the ADP-ribosyltransferase (gpAlt) of bacteriophage T4: ADP-ribosylation of E. coli RNA polymerase modulates T4 “early” transcription. Gene Expr. 1995;4(4-5):253–64.

66. Abedon ST. Look Who’s Talking: T-Even Phage Lysis Inhibition, the Granddaddy of Virus-Virus Intercellular Communication Research. Viruses. 2019;11(10).

67. Paddison P, Abedon ST, Dressman HK, Gailbreath K, Tracy J, Mosser E, et al. The roles of the bacteriophage T4 r genes in lysis inhibition and fine-structure genetics: a new perspective. Genetics. 1998;148(4):1539–50.

68. Agris PF, Vendeix FA, Graham WD. tRNA’s wobble decoding of the genome: 40 years of modification. J Mol Biol. 2007;366(1):1–13.

69. de Crecy-Lagard V, Hutinet G, Cediel-Becerra JDD, Yuan Y, Zallot R, Chevrette MG, et al. Biosynthesis and function of 7-deazaguanine derivatives in bacteria and phages. Microbiol Mol Biol Rev. 2024;88(1):e0019923.

70. Hutinet G, Kot W, Cui L, Hillebrand R, Balamkundu S, Gnanakalai S, et al. 7-Deazaguanine modifications protect phage DNA from host restriction systems. Nat Commun. 2019;10(1):5442.

71. Poisot T, Canard E, Mouquet N, Hochberg ME. A comparative study of ecological specialization estimators. Methods in Ecology and Evolution. 2012;3(3):537–44.

72. Bruto M, Labreuche Y, James A, Piel D, Chenivesse S, Petton B, et al. Ancestral gene acquisition as the key to virulence potential in environmental Vibrio populations. ISME J. 2018.

73. Corzett CH, Elsherbini J, Chien DM, Hehemann JH, Henschel A, Preheim SP, et al. Evolution of a Vegetarian Vibrio: Metabolic Specialization of Vibrio breoganii to Macroalgal Substrates. J Bacteriol. 2018;200(15).

74. Inoue T, Matsuzaki S, Tanaka S. A 26-kDa outer membrane protein, OmpK, common to Vibrio species is the receptor for a broad-host-range vibriophage, KVP40. FEMS Microbiol Lett. 1995;125(1):101–5.

75. Randall-Hazelbauer L, Schwartz M. Isolation of the bacteriophage lambda receptor from Escherichia coli. J Bacteriol. 1973;116(3):1436–46.

76. Roa M. Interaction of bacteriophage K10 with its receptor, the lamB protein of Escherichia coli. J Bacteriol. 1979;140(2):680–6.

77. Wandersman C, Schwartz M. Protein Ia and the lamB protein can replace each other in the constitution of an active receptor for the same coliphage. Proc Natl Acad Sci U S A. 1978;75(11):5636–9.

78. Castillo D, Rorbo N, Jorgensen J, Lange J, Tan D, Kalatzis PG, et al. Phage defense mechanisms and their genomic and phenotypic implications in the fish pathogen Vibrio anguillarum. FEMS Microbiol Ecol. 2019;95(3).

79. Nobrega FL, Vlot M, de Jonge PA, Dreesens LL, Beaumont HJE, Lavigne R, et al. Targeting mechanisms of tailed bacteriophages. Nat Rev Microbiol. 2018;16(12):760–73.

80. Maffei E, Shaidullina A, Burkolter M, Heyer Y, Estermann F, Druelle V, et al. Systematic exploration of Escherichia coli phage-host interactions with the BASEL phage collection. PLoS Biol. 2021;19(11):e3001424.

81. Pellizza L, Lopez JL, Vazquez S, Sycz G, Guimaraes BG, Rinaldi J, et al. Structure of the putative long tail fiber receptor-binding tip of a novel temperate bacteriophage from the Antarctic bacterium Bizionia argentinensis JUB59. J Struct Biol. 2020;212(1):107595.

82. Bartual SG, Otero JM, Garcia-Doval C, Llamas-Saiz AL, Kahn R, Fox GC, et al. Structure of the bacteriophage T4 long tail fiber receptor-binding tip. Proc Natl Acad Sci U S A. 2010;107(47):20287–92.

83. Schulz EC, Schwarzer D, Frank M, Stummeyer K, Muhlenhoff M, Dickmanns A, et al. Structural basis for the recognition and cleavage of polysialic acid by the bacteriophage K1F tailspike protein EndoNF. J Mol Biol. 2010;397(1):341–51.

84. Stummeyer K, Dickmanns A, Muhlenhoff M, Gerardy-Schahn R, Ficner R. Crystal structure of the polysialic acid-degrading endosialidase of bacteriophage K1F. Nat Struct Mol Biol. 2005;12(1):90–6.

85. Burnim AA, Dufault-Thompson K, Jiang X. The three-sided right-handed beta-helix is a versatile fold for glycan interactions. Glycobiology. 2024;34(7).

86. Keen EC. Tradeoffs in bacteriophage life histories. Bacteriophage. 2014;4(1):e28365.

87. Barcia-Cruz R, Goudenege D, Moura de Sousa JA, Piel D, Marbouty M, Rocha EPC, et al. Phage-inducible chromosomal minimalist islands (PICMIs), a novel family of small marine satellites of virulent phages. Nat Commun. 2024;15(1):664.

88. Cahier K, Piel D, Barcia-Cruz R, Goudenege D, Wegner KM, Monot M, et al. Environmental vibrio phage-bacteria interaction networks reflect the genetic structure of host populations. Environ Microbiol. 2023.

89. Bono LM, Draghi JA, Turner PE. Evolvability Costs of Niche Expansion. Trends Genet. 2020;36(1):14–23.

90. Al-Shayeb B, Sachdeva R, Chen LX, Ward F, Munk P, Devoto A, et al. Clades of huge phages from across Earth’s ecosystems. Nature. 2020;578(7795):425–31.

91. da Silva JD, Melo LDR, Santos SB, Kropinski AM, Xisto MF, Dias RS, et al. Genomic and proteomic characterization of vB_SauM-UFV_DC4, a novel Staphylococcus jumbo phage. Appl Microbiol Biotechnol. 2023;107(23):7231–50.

92. Hu M, Xing B, Yang M, Han R, Pan H, Guo H, et al. Characterization of a novel genus of jumbo phages and their application in wastewater treatment. iScience. 2023;26(6):106947.

93. Sharma R, Pielstick BA, Bell KA, Nieman TB, Stubbs OA, Yeates EL, et al. A Novel, Highly Related Jumbo Family of Bacteriophages That Were Isolated Against Erwinia. Front Microbiol. 2019;10:1533.

94. Edwards KF, Steward GF, Schvarcz CR. Making sense of virus size and the tradeoffs shaping viral fitness. Ecol Lett. 2021;24(2):363–73.

95. Rorbo N, Ronneseth A, Kalatzis PG, Rasmussen BB, Engell-Sorensen K, Kleppen HP, et al. Exploring the Effect of Phage Therapy in Preventing Vibrio anguillarum Infections in Cod and Turbot Larvae. Antibiotics (Basel). 2018;7(2).

96. Tan D, Dahl A, Middelboe M. Vibriophages Differentially Influence Biofilm Formation by Vibrio anguillarum Strains. Appl Environ Microbiol. 2015;81(13):4489–97.

97. Tan D, Svenningsen SL, Middelboe M. Quorum Sensing Determines the Choice of Antiphage Defense Strategy in Vibrio anguillarum. mBio. 2015;6(3):e00627.

