## Supplementary Figures for "Generalists in a Specialist World: Vibrio-Phages with Broad Host Range"

**SUPPLEMENTARY INFORMATION**

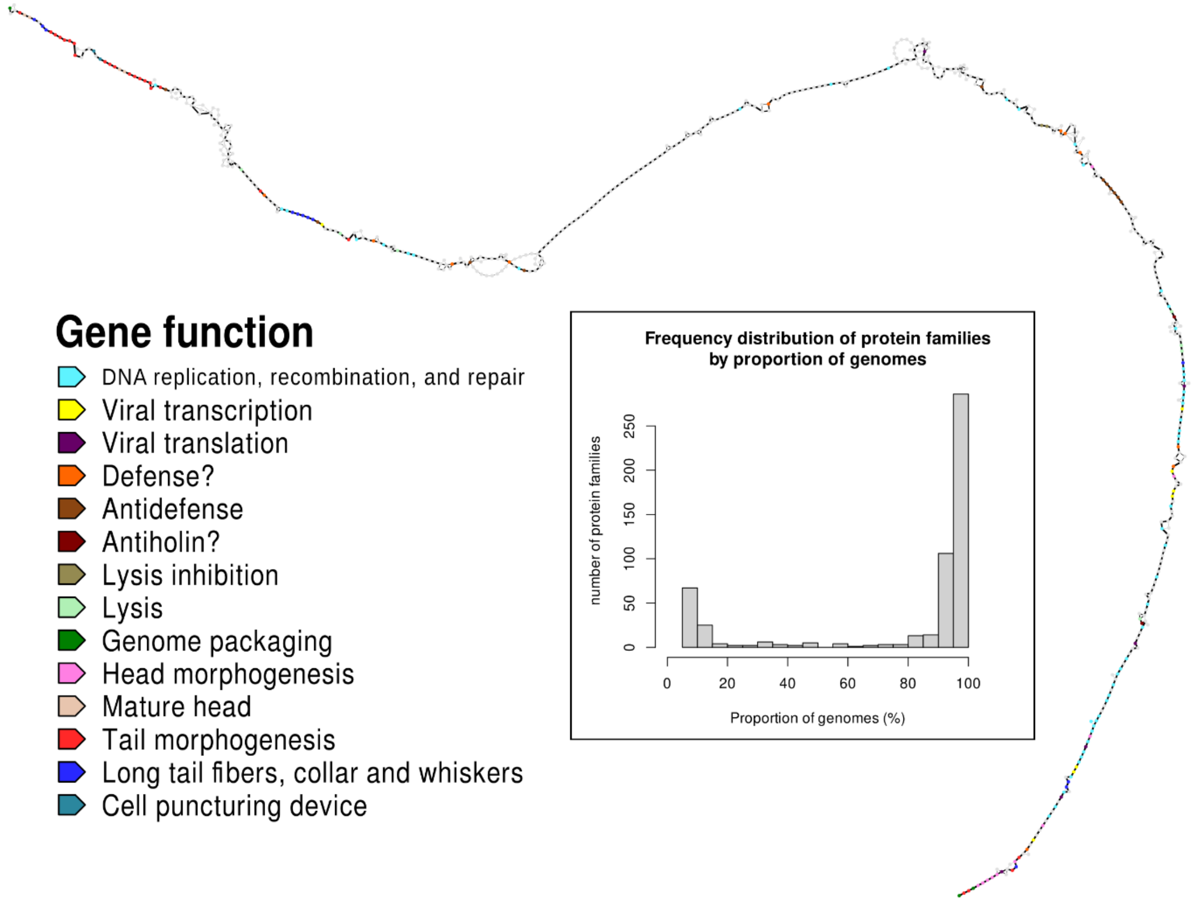


**Supplementary Figure 1**. **The pangenome of the Schizotequatrovirus**. The histogram illustrates the number of protein families (groups of BBHs) as a function of the proportion of encoding genomes. Thus, families on the left of the histogram represent accessory genes, while those on the right correspond to core genes. Surrounding the histogram is a network that depicts the genomic organization of the pangenome, where nodes represent BBH groups and edges connect BBH groups when their members are found adjacent to each other in at least one phage genome. The color of each BBH group corresponds to its functional category, as indicated by the legend on the left, with uncharacterized genes shown in grey. The width of each edge is proportional to the frequency of adjacency between two BBH groups across the 18 phage genomes.

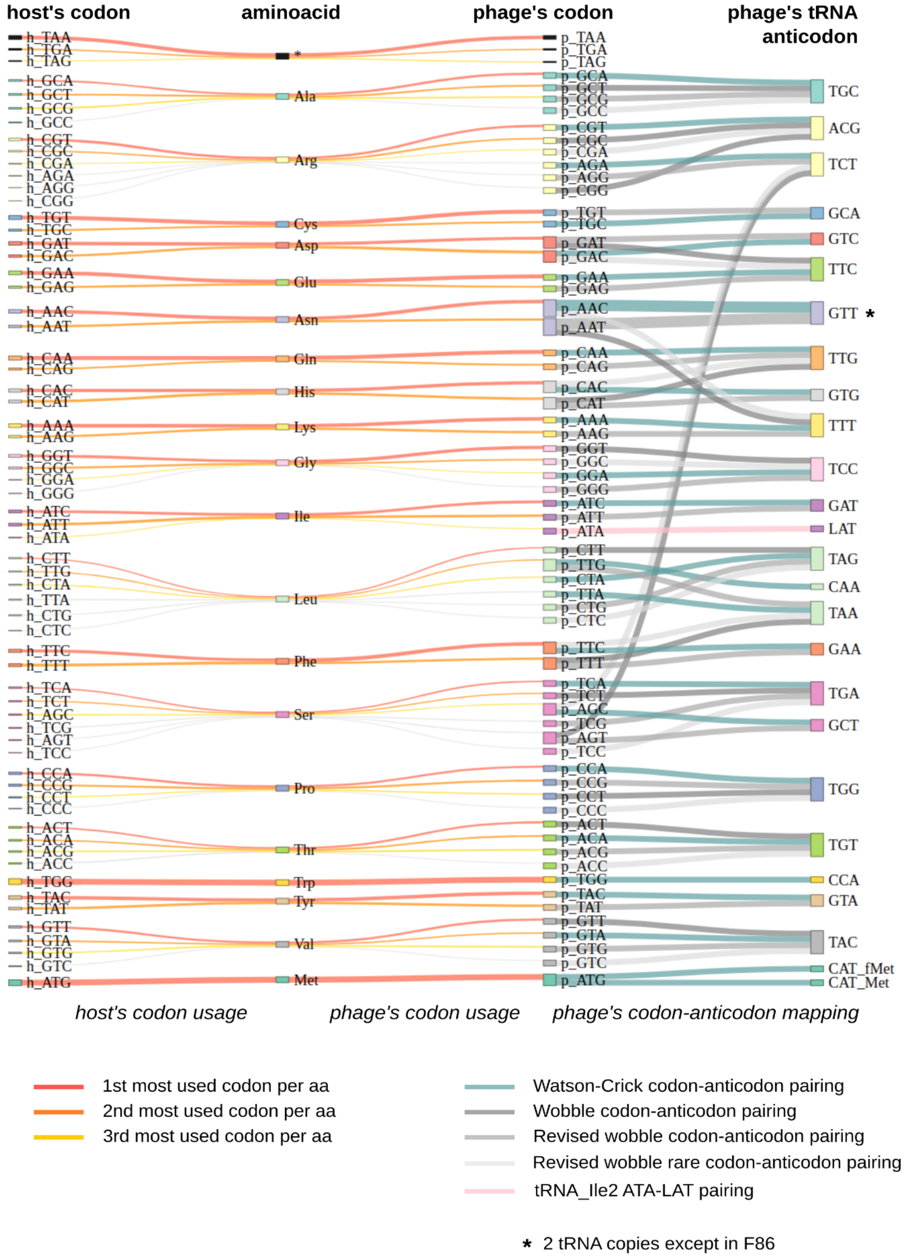


**Supplementary Figure 2. tRNAs of the Schizotequatrovirus.** The four node levels of the Sankey multipartite network represent, in order: host codons, amino acids, phage codons, and the 26 distinct anticodons of the phage tRNAs. Edges connect codons to their corresponding amino acid and to their associated tRNA anticodons. tRNA-fMet-CAT anticodon pairs with starting ATG codons while tRNA-Met-CAT anticodon pairs with standard ATG codons. The width of each edge reflects the host codon usage, the phage codon usage, and the average number of tRNA copies among the 18 phage genomes (1 copy in all genomes except for tRNA-Asn-GTT which is present in one copy in F86 and in two copies in the 17 other phages). The links between codons and amino acids are colored according to the codon usage hierarchy for each amino acid (see legend at the bottom left). Codon-anticodon links are colored based on the type of pairing: blue for perfect Watson-Crick pairing, pink for the post-transcriptionally modified LAT anticodon of tRNA-Ile2-CAT pairing with the ATA codon, and grey for Wobble pairing.

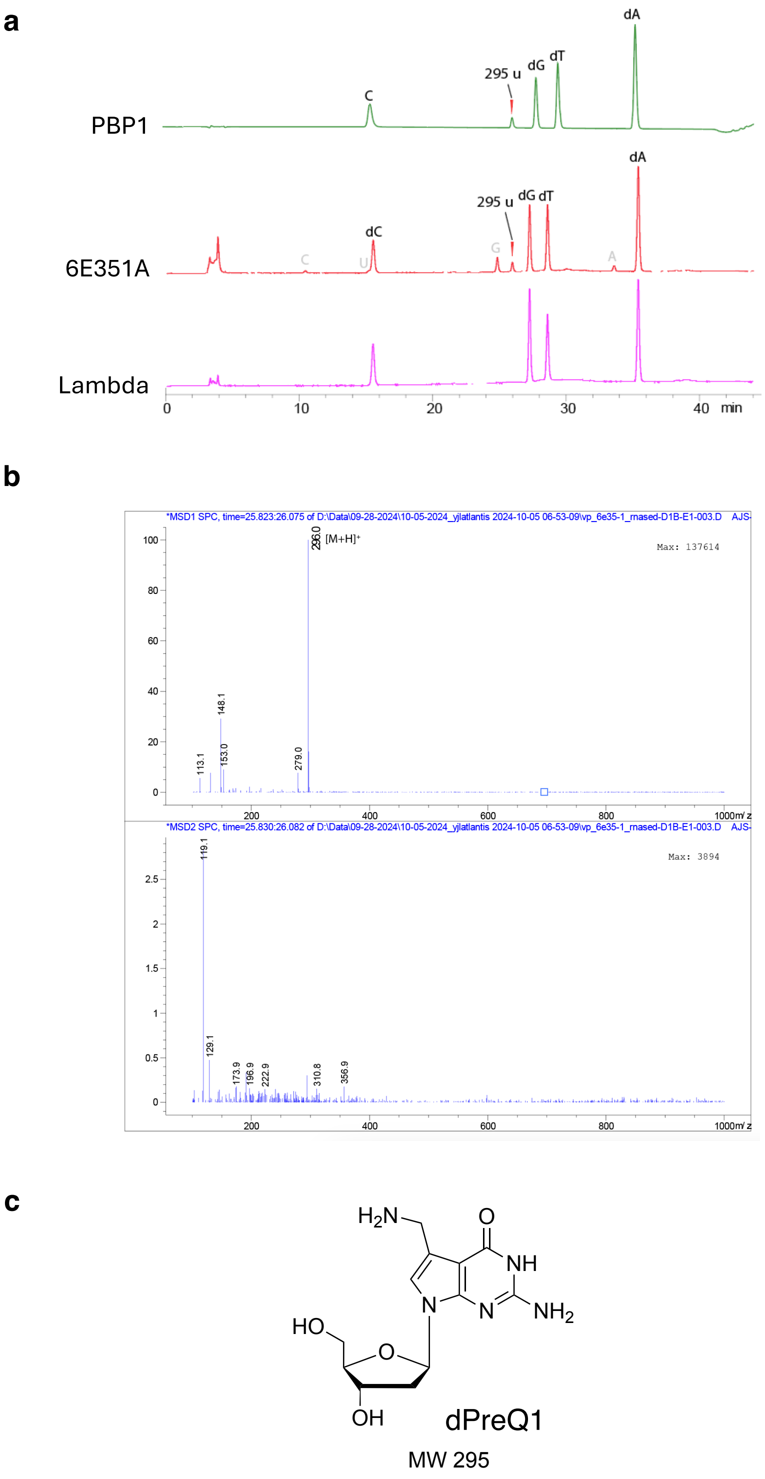


**Supplementary Figure 3. HPLC-MS analysis of dG modification in 6E351A DNA revealing dPreQ1 (7-deaza-dG) modification.** The 6E351A DNA contains 15% dG modification and has an observed mass of 295 Da, which matches the *dPreQ1*, a 7-deaza-dG modification. **a** About 0.5 µg of RNase-treated, purified DNA (phage PBP1, 6E351A or Lambda) was hydrolyzed to nucleoside (NEB #M0649) and subjected nucleoside solution to HPLC-MS analysis. **b** MSD signal at 25.8 min corresponds to a MW of 295. **c** This mass matches the dPreQ1, a 7-deaza-dG modification.

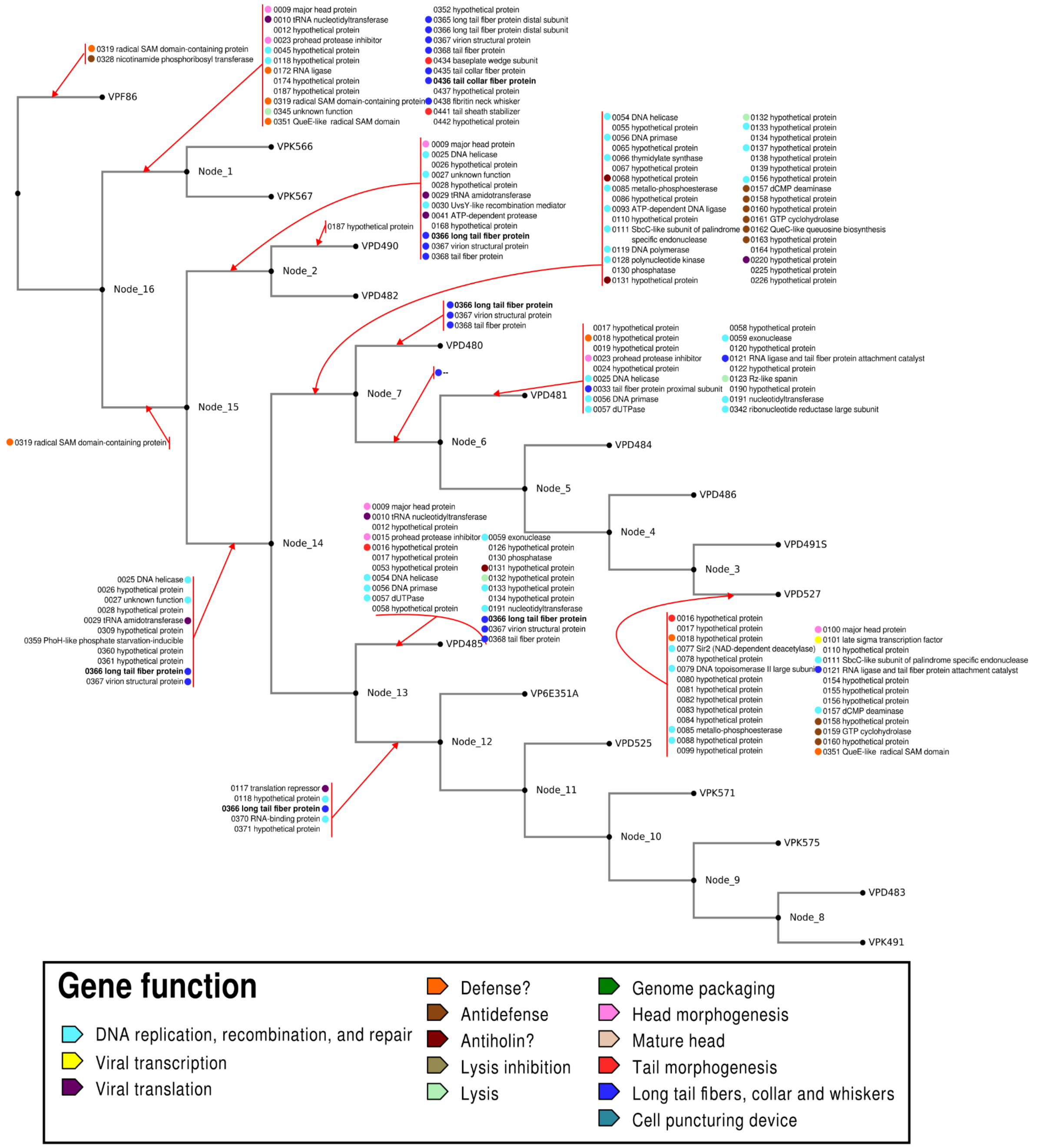


**Supplementary Figure 4. Main events of homologous recombination.** Each red arrow indicates a branch of the recombination-free phylogeny on which a main event of recombination in core genes is inferred by Gubbins to have occurred (negative log likelihood > 500). All groups of recombinant core BBHs are labeled based on the annotation of the orthologs in 6E351A. Colored circles indicate the functional category of core BBH groups.

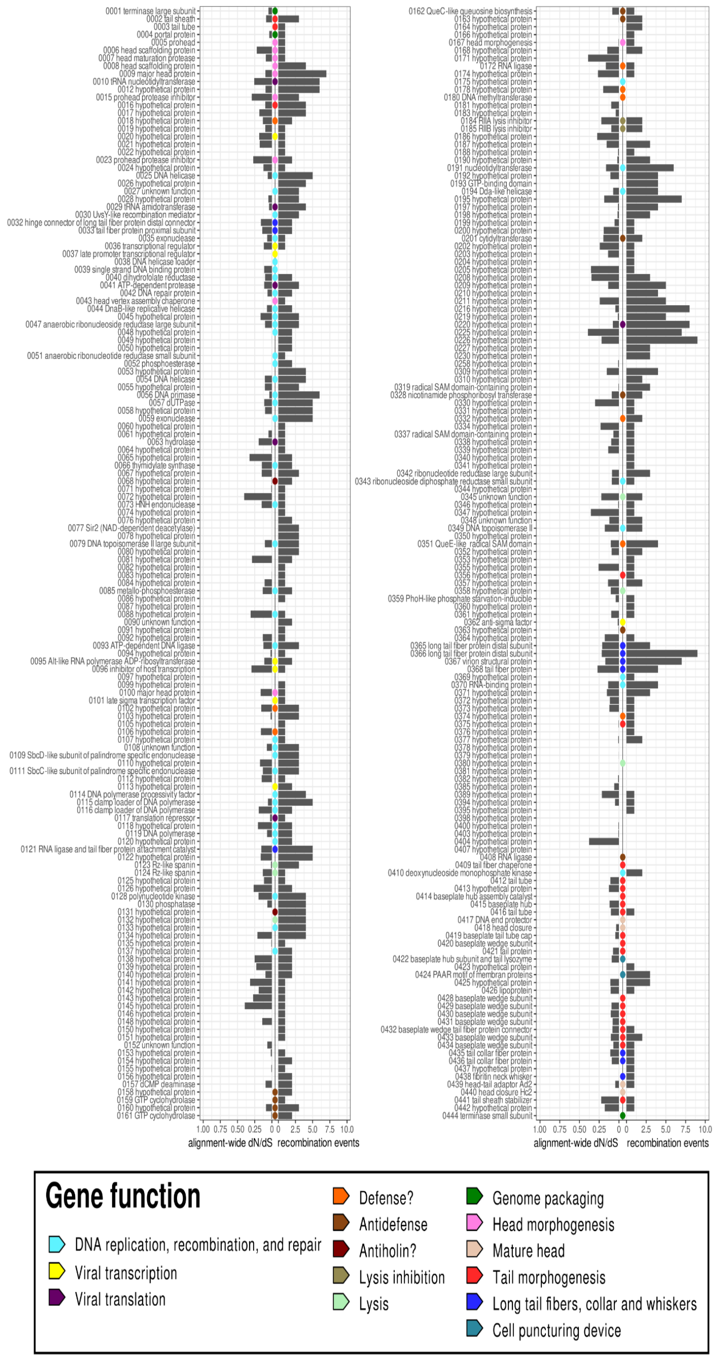


**Supplementary Figure 5**. **Number of recombination events and strength of purifying selection at each core locus.** The lines represent core gene loci present in all 18 phage genomes, and ordered according to their position in the reference genome 6E351A, with gene labels corresponding to functional annotations provided by Pharokka in 6E351A. At the center of each line, a colored circle indicates the functional category of the core gene (legend at the bottom); genes without circles being of uncharacterized function. Two key metrics are shown for each persistent locus. The first is the alignment-wide dN/dS (the ratio of nonsynonymous to synonymous mutation fixation rates), which estimates the strength of purifying selection acting on the gene. A dN/dS ratio below 1 suggests purifying selection, with values closer to 0 indicating stronger selective pressure against amino-acid substitutions. The second value is the number of recombination events inferred by Gubbins to have occurred at the locus along the phage recombination-free phylogeny. Regions with consecutive core loci exhibiting high recombination event densities can be identified as recombination hotspots.

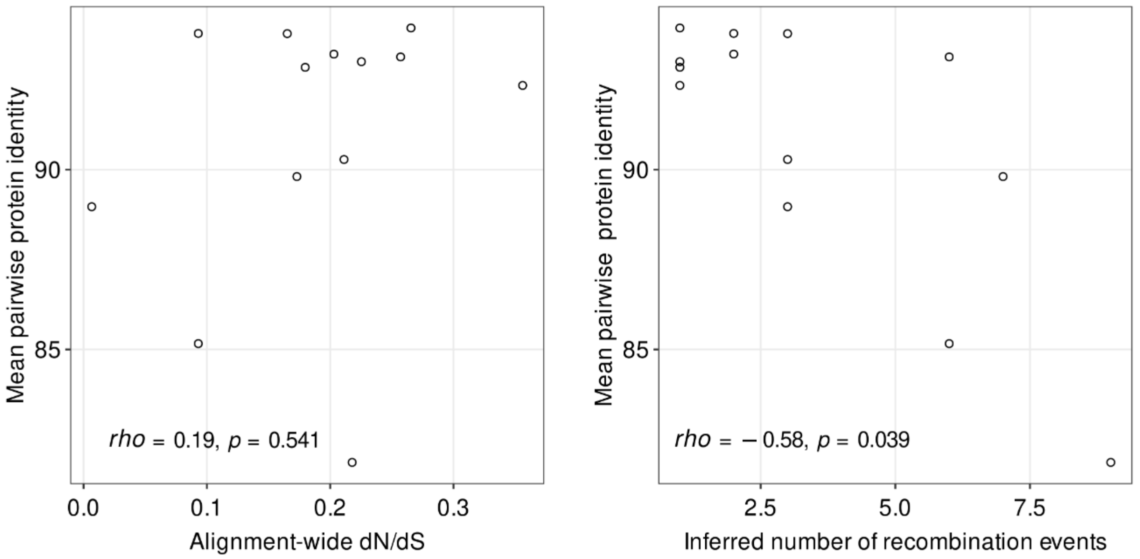


**Supplementary Figure 6. Protein-level divergence within core divergent BBH groups is associated with the number of inferred recombination events, not with the estimated alignment-wide dN/dS ratio.** Each dot represents a core BBH group yielding a mean pairwise identity <95% in the clade of the 17 closely-related phages. The y-axis shows the mean pairwise protein sequence identity between BBHs from the same core BBH group. The x axis gives either the dN/dS rate (left) or the number of recombination events (right) within the BBH group. Spearman correlation statistics are displayed at the bottom of each scatter plot.

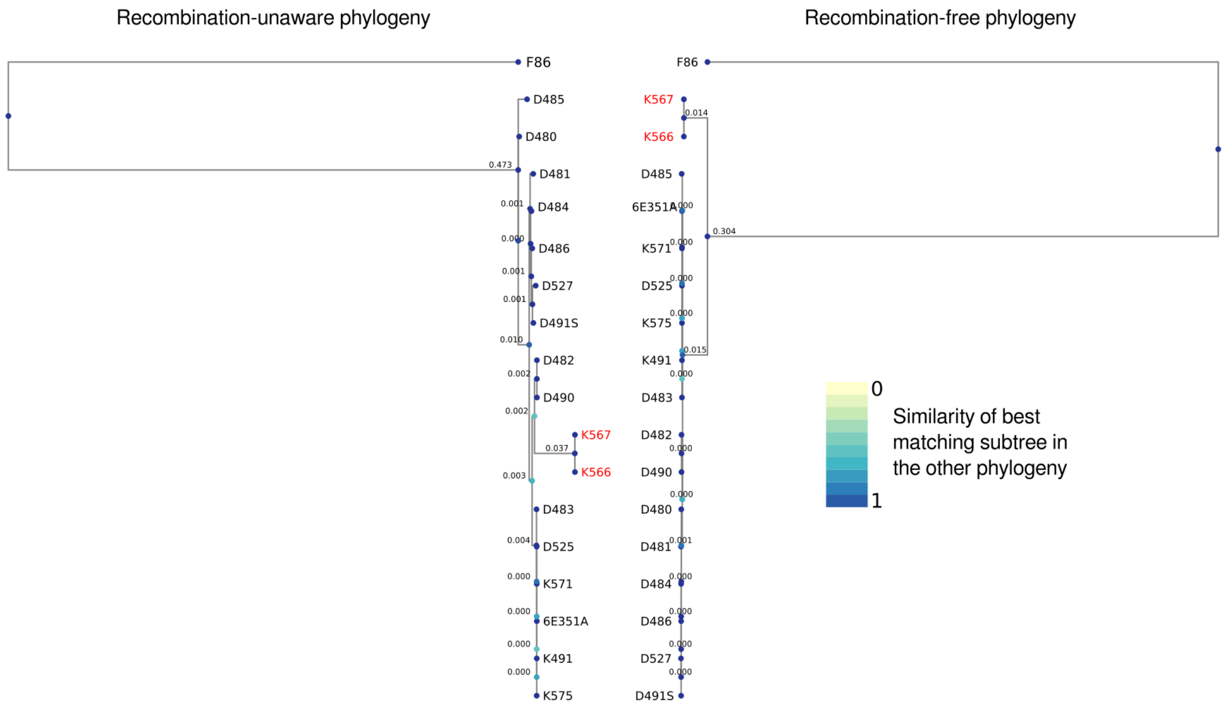


**Supplementary Figure 7. Recombination-unaware vs recombination-free phylogeny of Schizotequatroviruses**. The estimated average nucleotide substitution per site is written for each internal branch. The color of each node of a phylogeny highlights the similarity of the best matching subtree in the other phylogeny. The clade formed by phages K566 and 567 is highlighted in red on both trees.

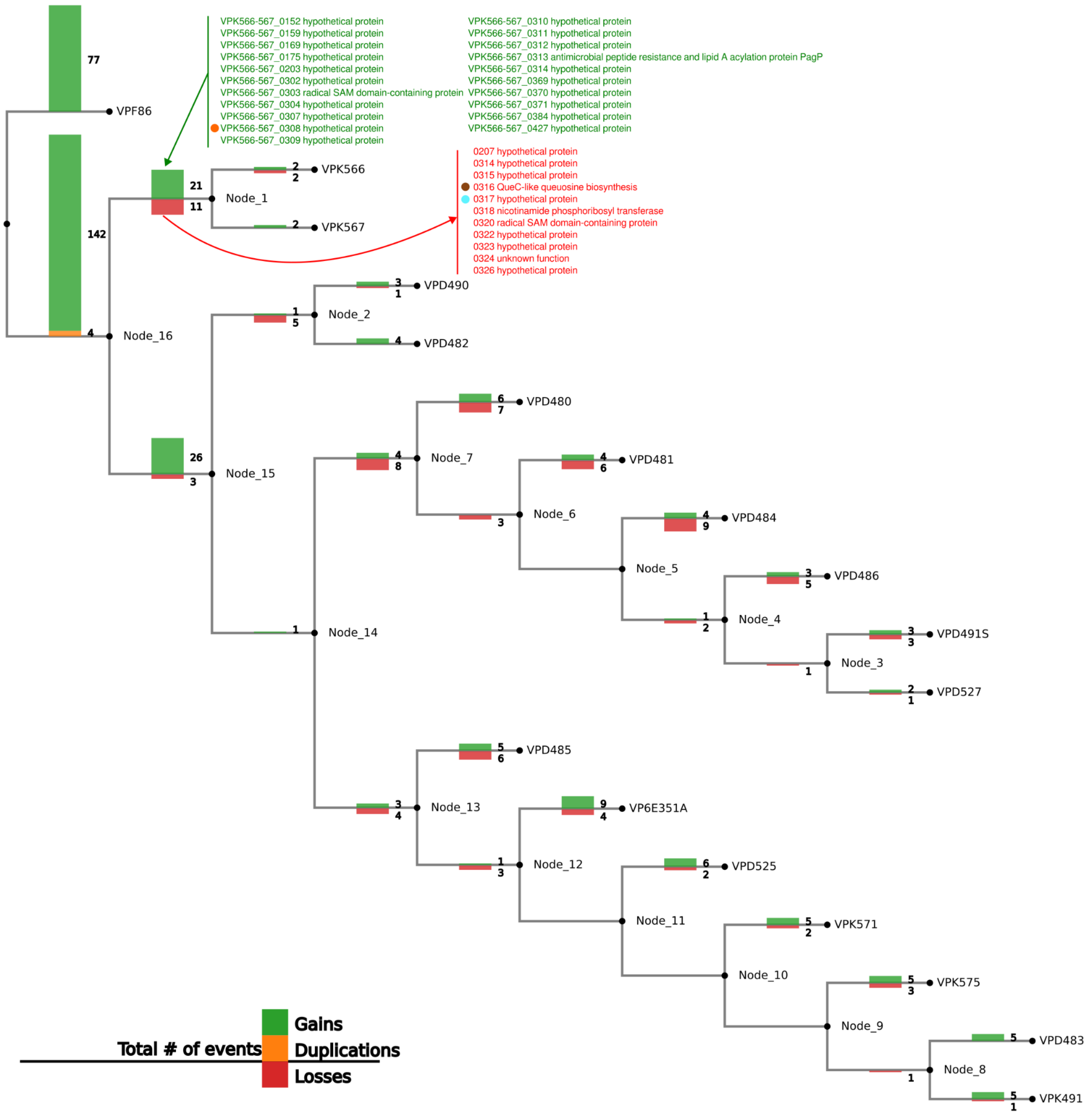


**Supplementary Figure 8. Phylostratigraphy.** The stacked histogram on each branch of the recombination-free phylogeny indicates the number of inferred events of gains, losses or duplications of gene families (identified as Hierarchical Orthologous Groups by the OMA orthology inference pipeline) relative to the parent branch. The red arrow indicates the ancestral protein families (annotation of 6E351A) inferred to have been lost in the last common ancestor of phages K566 and K567, while the green arrow indicates the families inferred to have been gained (annotation of K566). An entire ancestral island (0314 to 0326 in 6E351A) is predicted to have been replaced by another one (0302 to 0314 in K566) in this ancestor.

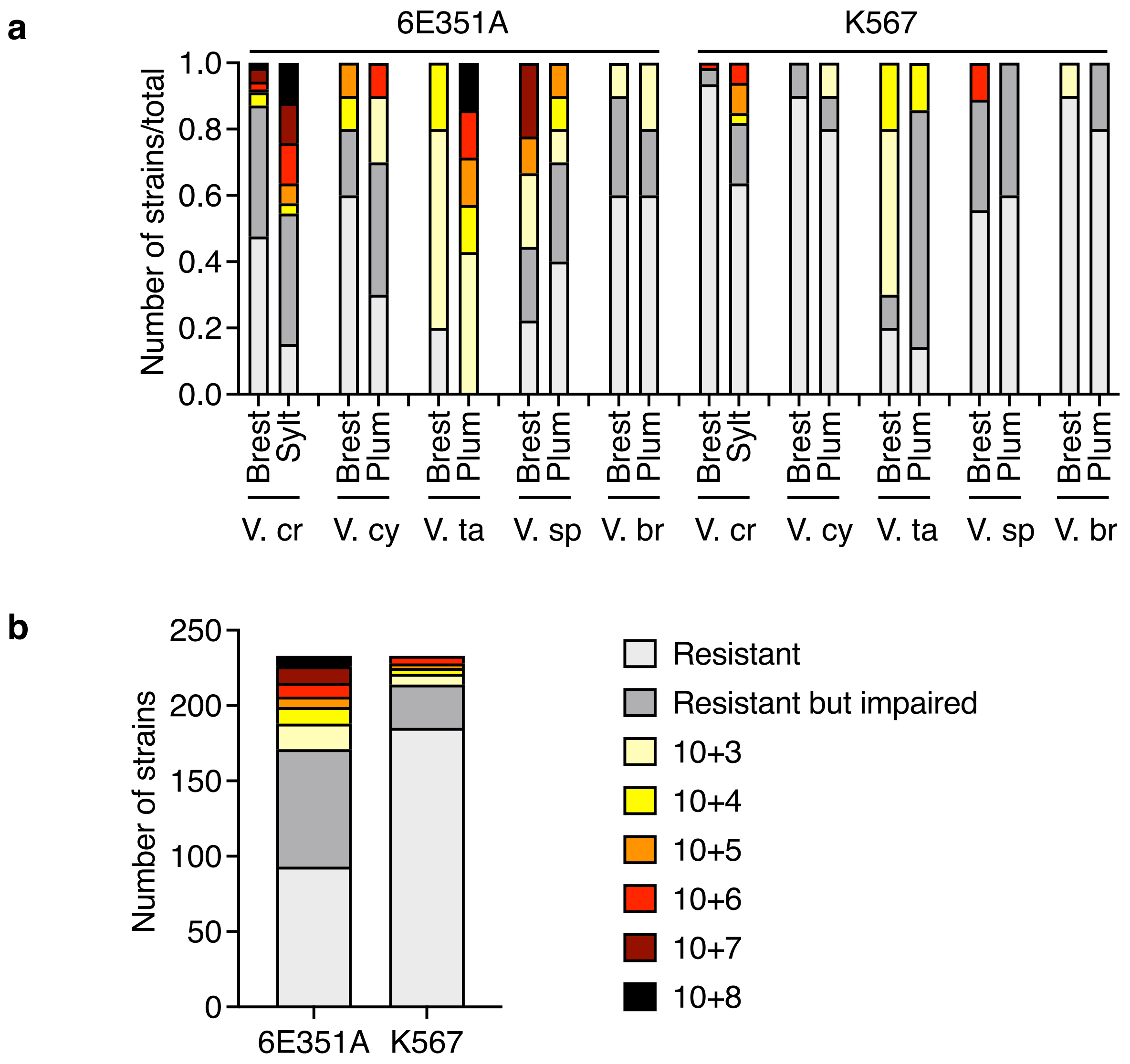


**Supplementary Figure 9. Host Range and Sensitivity of Phages 6E351A and K567 Across *Vibrio* Species and Geographic Locations.** The titer of phages 6E351A and K567, isolated in Brest (France), was estimated on members of diverse species (*V. crassostreae*, V cr; *V. cyclitrophicus*, V. cy; *V. tasmaniensis*, V. ta; *V. splendidus*, V. sp; *V. breoganii,* V. br) isolated at the same or different locations (Sylt, Germany; Plum Island, USA). Hosts were classified as sensitive when the phage titers ranged from $10^3$ to $10^8$ PFU /mL, resistant but impaired, when we observed a clearing zone but no production of viable phages and resistant when no plaque, nor clearing zone was observed. **a** Bar charts indicate the number of strains on the total number of strains for each category per location and species (124 and 33 strains from Brest and Sylt respectively *for V. crassostreae*; 10 strains per species and location for the other species). **b** Bar charts indicate the number of strains for each category per infecting phage.

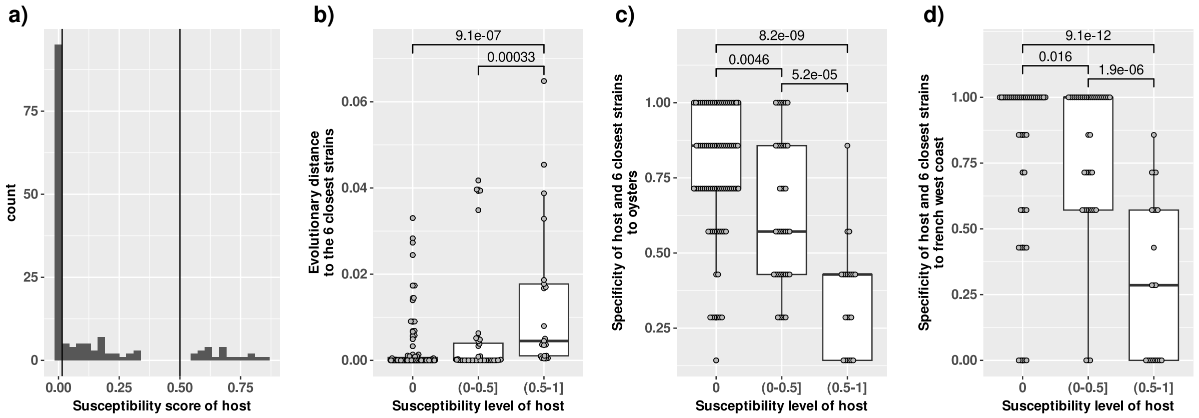


**Supplementary Figure 10**. **Resistant hosts are in clades characterized with a low degree of polymorphism in core genes and a certain specificity to oysters and to the french West coast**. The panel **a** displays the distribution of susceptibility indexes among the host population. The distribution clearly delineates three classes of susceptibility indexes: 0 (resistant bacteria), above 0 to 0.5 (slightly susceptible bacteria), and above to 0.5 to 1 (highly susceptible bacteria). The panel **b** shows the distribution of the degree of polymorphism in core genes to the 6 closest strains of the phylogeny for each susceptibility class (given as the mean of the 6 pairwise patristic distances in the host phylogeny). The panel **c** shows the distribution of the specificity of hosts and their 6 closest strains to the oyster biome for each susceptibility class. The panel **d** shows the distribution of the specificity of hosts and their 6 closest strains to the french West coast (Brest) for each susceptibility class. Significant differences in distributions between classes are given by p-values of Mann-Whitney tests. Each boxplot gives the median, the first and third quartiles (the two hinges). The upper whisker extends from the hinge to the largest value no further than 1.5 * IQR from the hinge (where IQR is the inter-quartile range, or distance between the first and third quartiles) and the lower whisker extends from the hinge to the smallest value at most 1.5 * IQR of the hinge.

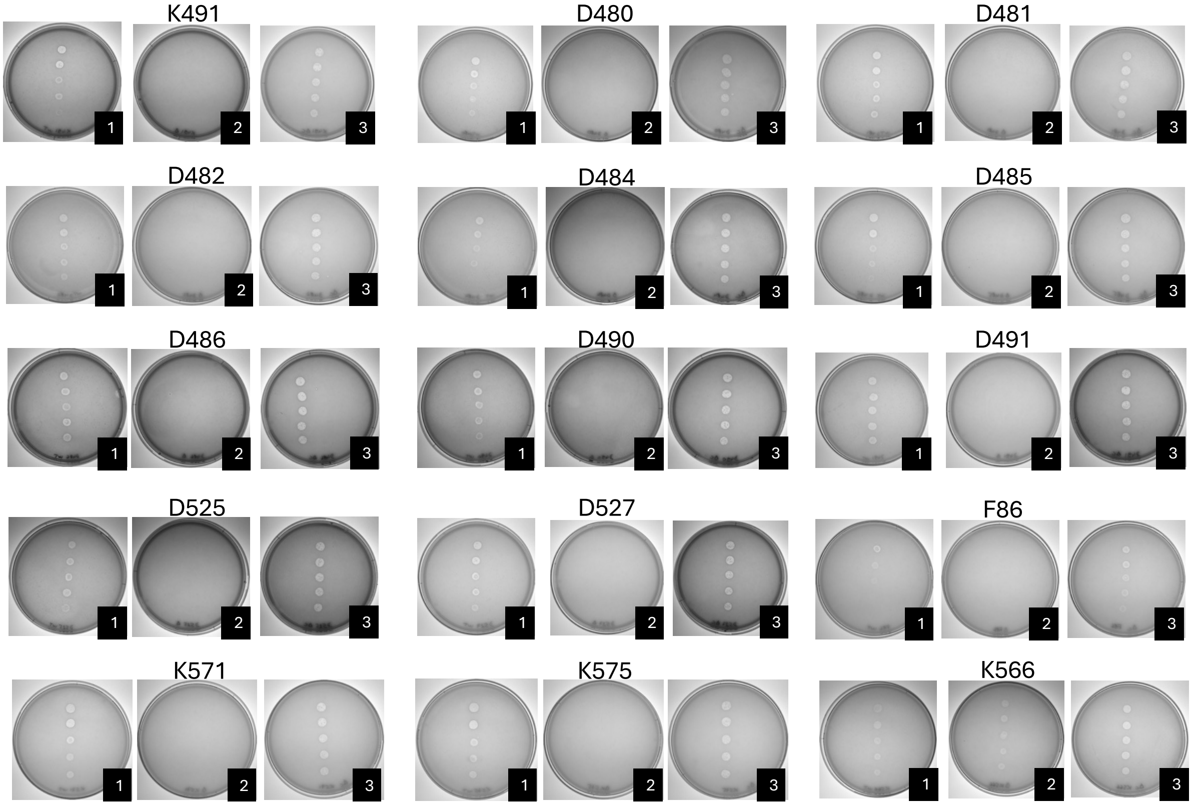


**Supplementary Figure 11. Role of OmpK as a Receptor for Schizotequatroviruses.** A killing assay was performed by spotting 10-fold serial dilutions of the phage on the wild-type strain 22_O_6 (1), its Δ*ompK* mutant derivative (2), and the Δ*ompK* strain complemented with a plasmid constitutively expressing *ompK* (3).

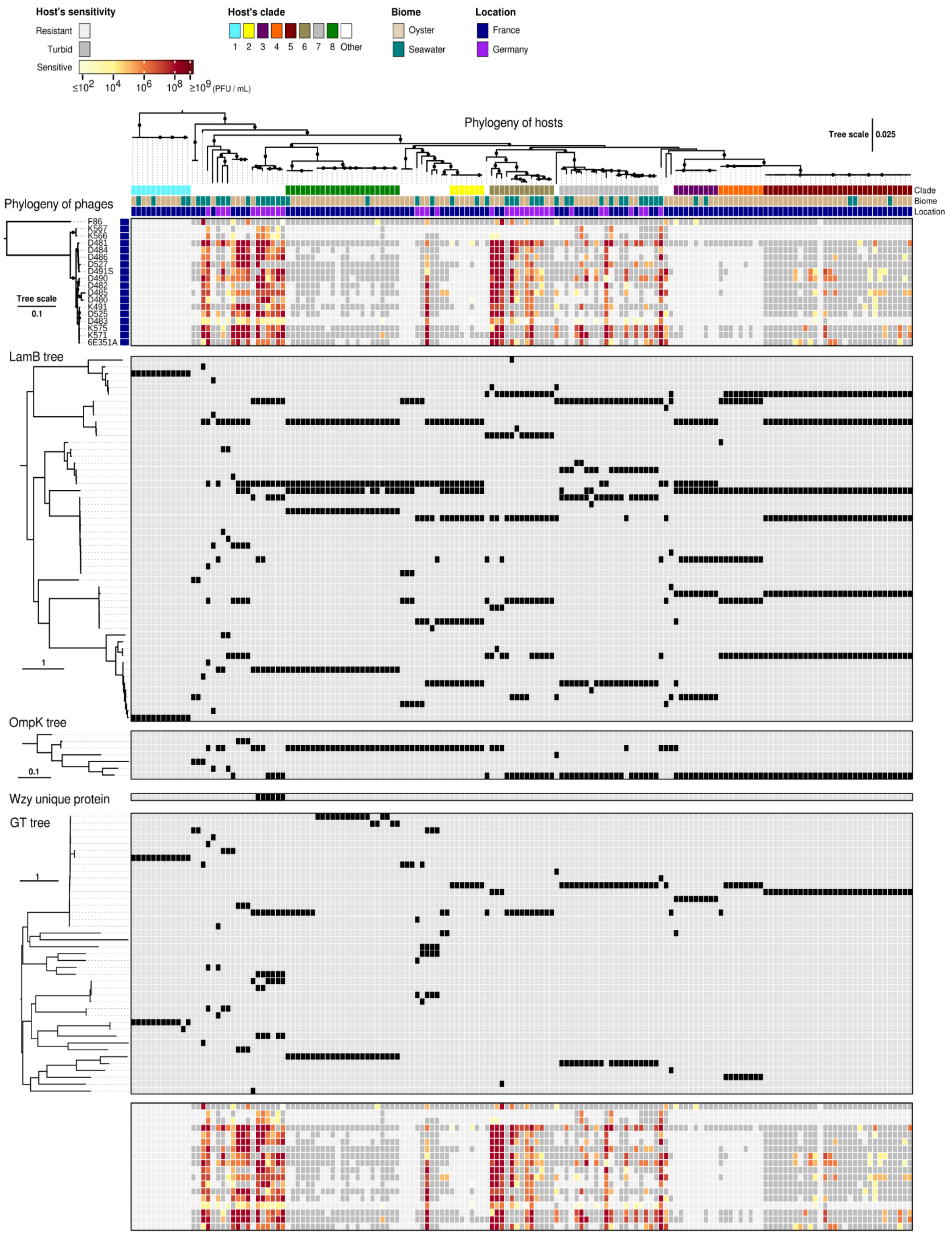


**Supplementary Figure 12. Phylogenetic profiling of LamB, OmpK, Wzy and GT alleles.** The heatmaps at the top and the bottom correspond to the phages-bacteria interaction matrix. The trees on the left correspond to the mid-point rooted phylogenies of unique variants among the LamB, OmpK, Wzy and GT families. The matrix on the right of each tree gives the presence/absence of each variant across host genomes. Note that the OmpK gene is absent in resistant hosts from clade V1. Phages K566 and K567 use LamB for adsorption but no LamB variant is specific to hosts that these phages can infect. Similarly, the presence of EpsG family protein (Wzy) or the phylogeny of glycosyltransferase (GT) does not fully explain host range, even though these proteins facilitate adsorption.

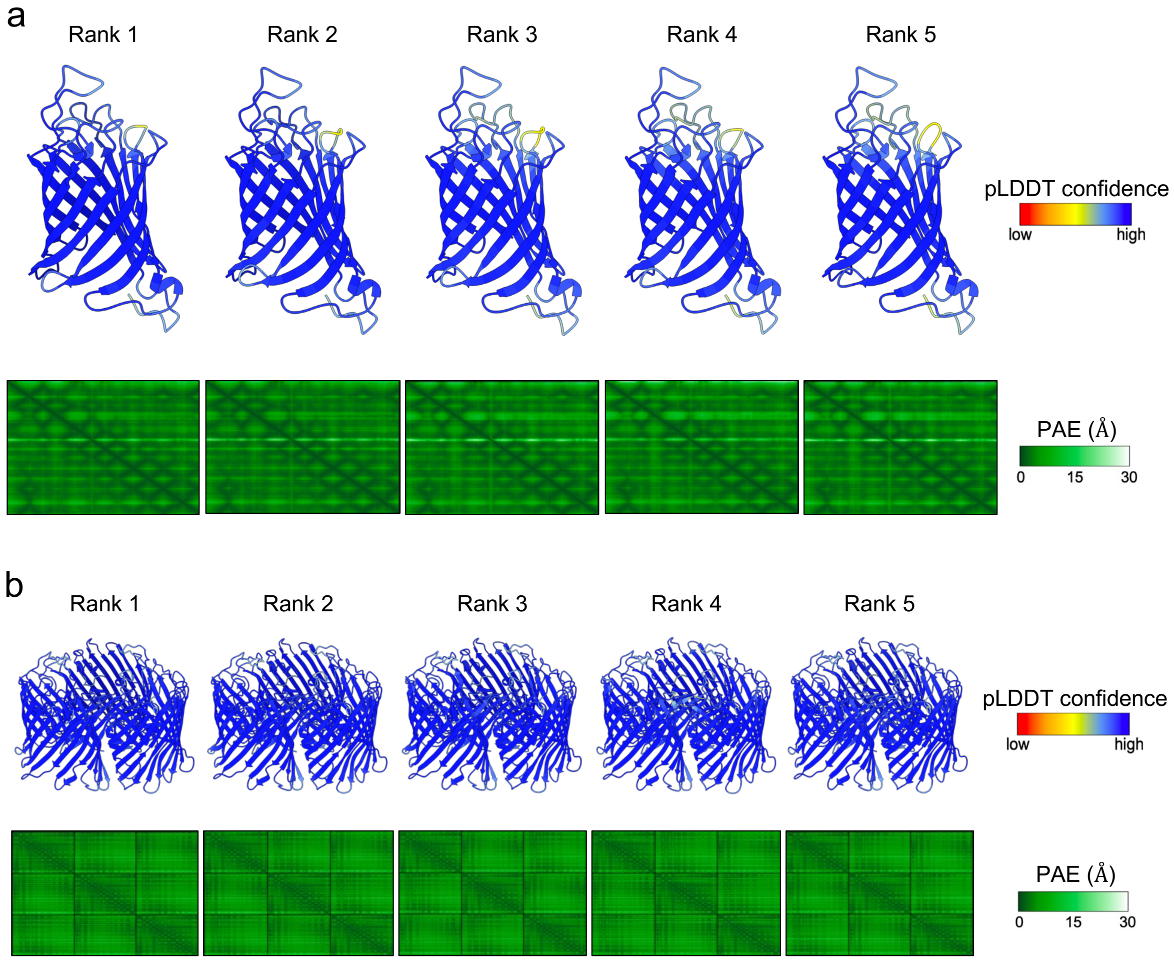


**Supplementary Figure 13 a** OmpK structures, with the predicted signal sequence (residues 1-20) removed, predicted by AlphaFold2 as implemented within ColabFold. **b** LamB trimer structures, with the predicted signal sequence (residues 1-22) removed, predicted by AlphaFold-multimer as implemented within ColabFold. Structures are aligned and shown from the same orientation. Structures are ranked according to the predicted template modeling (pTM) score and are colored according to the predicted local distance difference test (pLDDT) score, which indicates per-residue model confidence for the individual protein chains within the complex. Confidence in the prediction of the complex is indicated by the predicted aligned error (PAE) scores, which indicate positional error in angstroms for a given pair of residues across all protein chains.

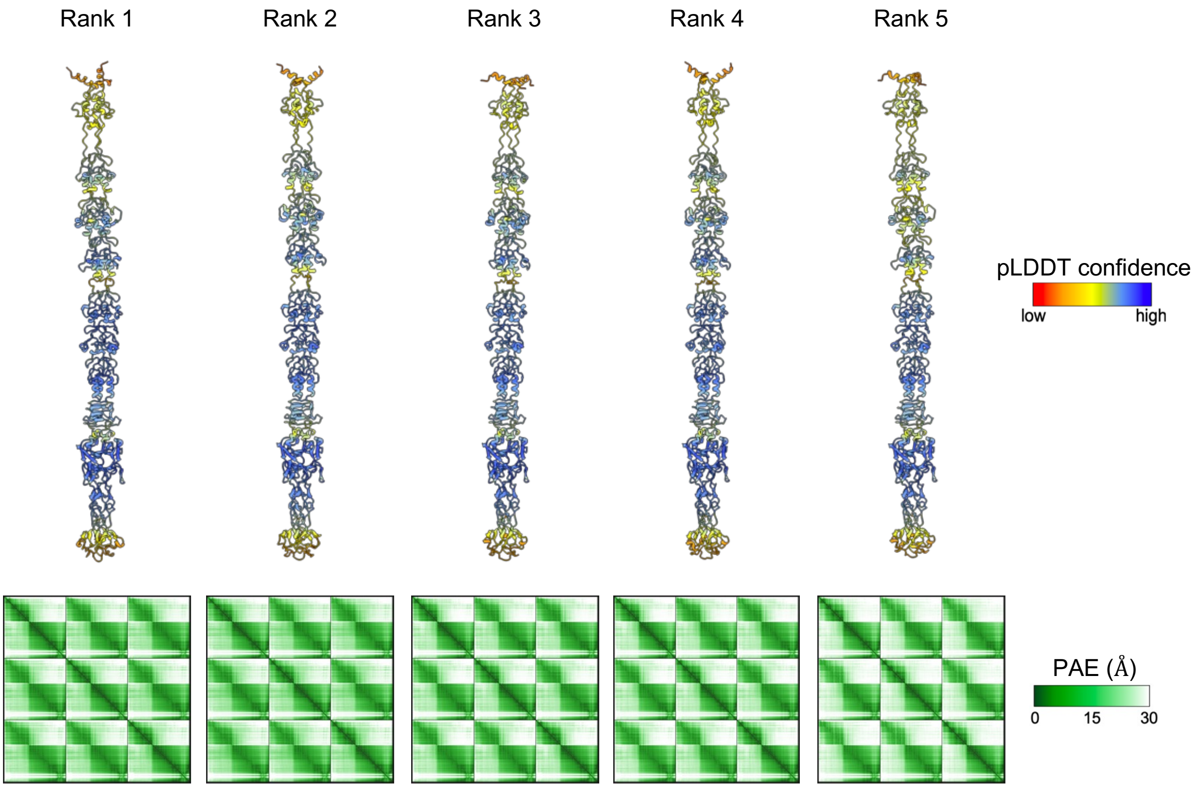


**Supplementary Figure 14** Trimeric structures of VPK566_0422 in complex with two $Mg^{2+}$ ions predicted by AlphaFold3. Structures are aligned and shown from the same orientation. Structures are ranked according to the predicted template modeling (pTM) score and are colored according to the predicted local distance difference test (pLDDT) score, which indicates per-residue model confidence for the individual protein chains within the complex. Confidence in the prediction of the complex is indicated by the predicted aligned error (PAE) scores, which indicate positional error in angstroms for a given pair of residues across all protein chains.

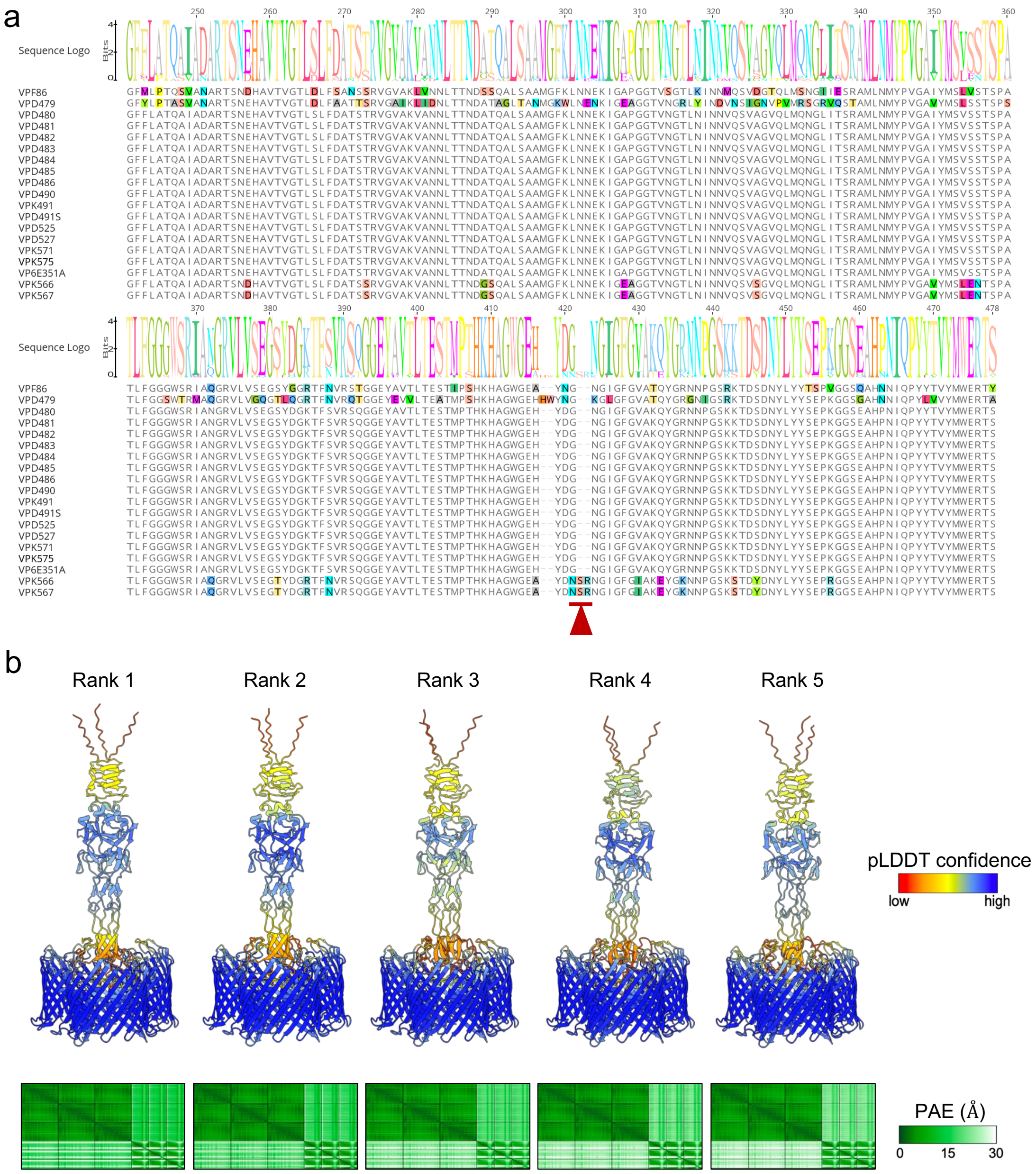


**Supplementary Figure 15 a** Multiple sequence alignment of the C-terminal half of VP6E351A_0436 and its orthologs from 18 additional *Vibrio* phages (VPs). The consensus sequence is indicated at the top of the alignment as a sequence logo. Deviations from the consensus are highlighted. The phage K566/K567-specific sequence variation of interest is highlighted by the red arrow. **b** Structures of trimeric LamB, with the predicted signal sequence (residues 1-22) removed, in complex with residues 295-476 of trimeric VPK566_0422, predicted by AlphaFold-multimer as implemented within ColabFold. Structures are aligned and shown from the same orientation. Structures are ranked according to the predicted template modeling (pTM) score and are colored according to the predicted local distance difference test (pLDDT) score, which indicates per-residue model confidence for the individual protein chains within the complex. Confidence in the prediction of the complex is indicated by the predicted aligned error (PAE) scores, which indicate positional error in angstroms for a given pair of residues across all protein chains.

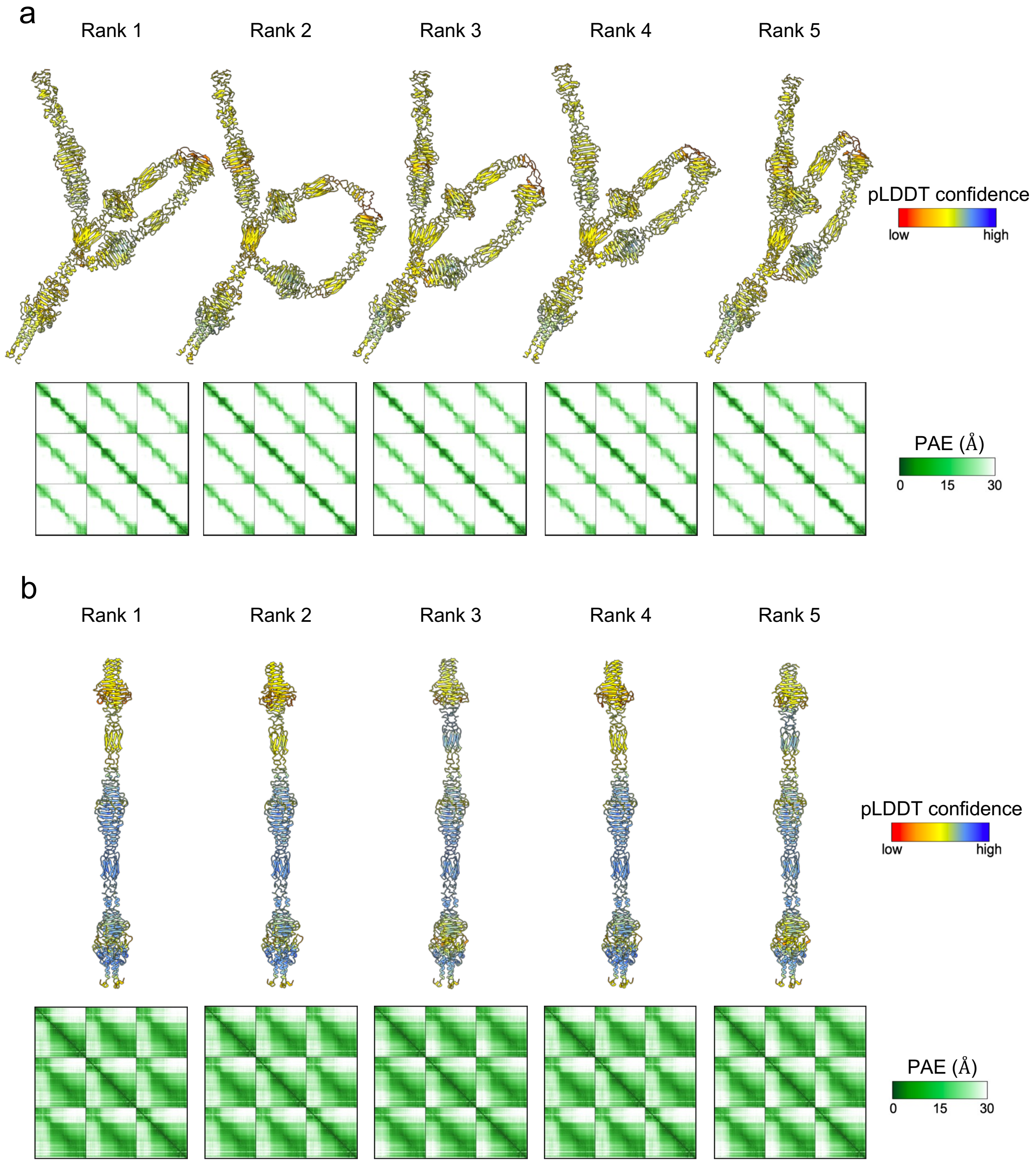


**Supplementary Figure 16** Trimeric structures of **a** VP6E351A_0365 and **b** VP6E351A_0366 predicted by AlphaFold3. Structures are aligned and shown from the same orientation. Structures are ranked according to the predicted template modeling (pTM) score and are colored according to the predicted local distance difference test (pLDDT) score, which indicates per-residue model confidence for the individual protein chains within the complex. Confidence in the prediction of the complex is indicated by the predicted aligned error (PAE) scores, which indicate positional error in angstroms for a given pair of residues across all protein chains.

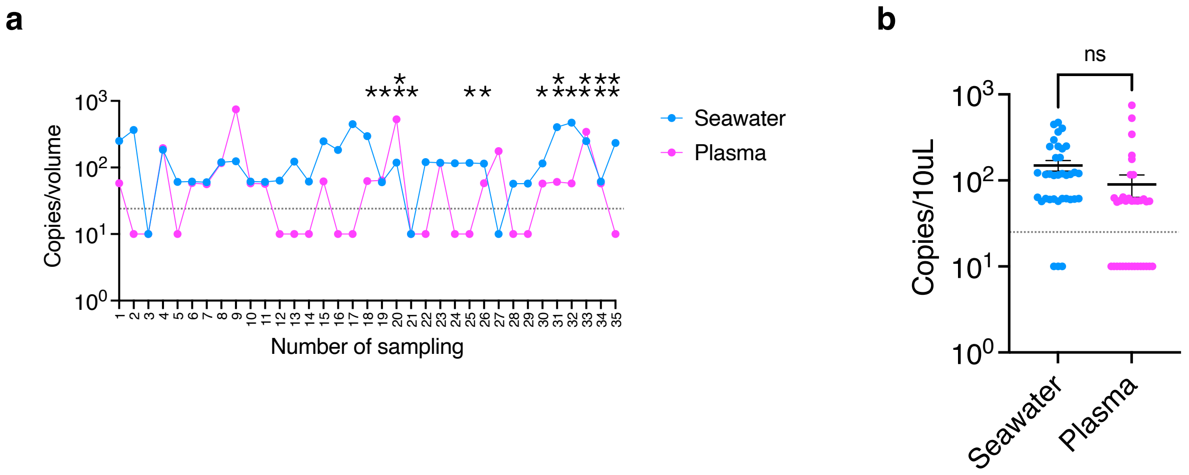


**Supplementary Figure 17 Absolute abundance of Schizotequatroviruses in seawater and oysters. a** The number of genome copies per Liter of seawater or mL of oyster plasma (pooled from 90 oysters) was quantified using droplet digital PCR (ddPCR). DNA samples were prepared from virome sources collected over 35 sampling dates during the 2021 time series. Asterisks indicate the sampling dates when one (*) or two (**) phages were isolated. The dashed line represents the detection limit. **b** A paired t-test revealed no significant difference in the abundance of Schizotequatroviruses between seawater and oyster plasma. Bars represent the mean ± SEM.

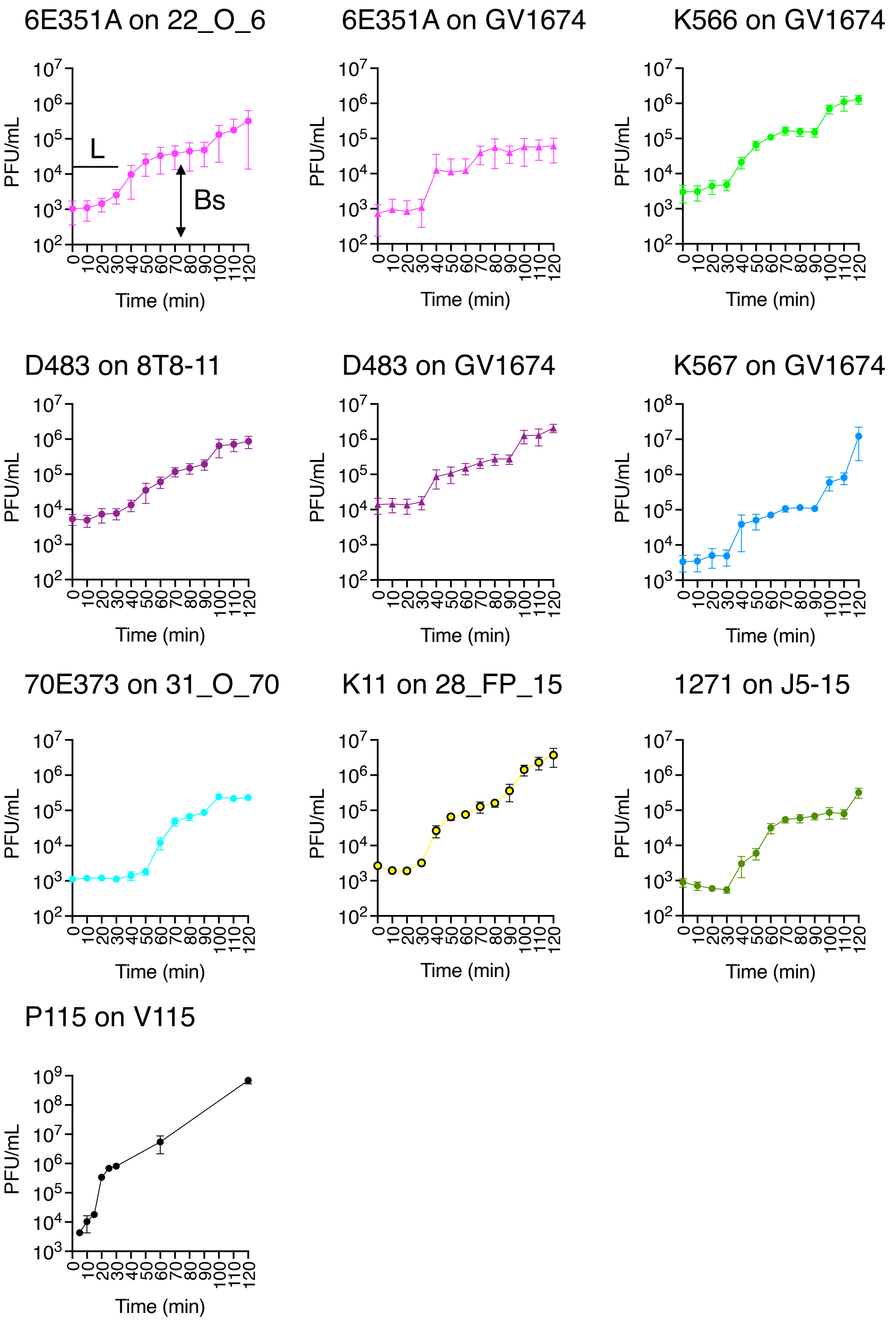


**Supplementary Figure 18 One-step growth curves.** Schizotequatroviruses (6E351A, D483, K566, and K567) were tested on their original host (circles) or an alternative host (triangles). For comparison, representative members of genera P1_12 (70E373), P2_4 (K11), and P8_8 (1271), which infect *V. crassostreae* from clades V1, V2, and V8, respectively, as well as phage P115, highly specific to a strain of *V. chagasii,* were also analyzed. Bars represent the mean ± standard error of the mean (SEM) from four independent replicates. L: latency period; Bs: burst size.
